# OMA1 mediates local and global stress responses against protein misfolding in CHCHD10 mitochondrial myopathy

**DOI:** 10.1101/2021.12.21.473493

**Authors:** Mario K. Shammas, Xiaoping Huang, Beverly P. Wu, Insung Song, Nicholas Randolph, Yan Li, Christopher K. E. Bleck, Danielle A. Springer, Carl Fratter, Ines A. Barbosa, Andrew F. Powers, Pedro M. Quirós, Carlos Lopez-Otin, Joanna Poulton, Derek P. Narendra

**Affiliations:** Inherited Movement Disorders Unit, Neurogenetics Branch, National Institute of Neurological Disorders and Stroke, National Institutes of Health, Bethesda, MD 20892, USA.; Protein/peptide sequencing facility, National Institute of Neurological Disorders and Stroke, National Institutes of Health, Bethesda, MD 20892, USA; National Heart, Lung, and Blood Institute, National Institutes of Health, 9000 Rockville Pike, Bethesda, MD, 20892, USA; Oxford Genetics Laboratories, Oxford University Hospitals NHS Foundation Trust, Oxford, UK; Department of Medical and Molecular Genetics, School of Basic & Medical Biosciences, King’s College London, UK; Ionis Pharmaceuticals, Carlsbad, CA 92010, USA; Departamento de Bioquímica y Biología Molecular, Facultad de Medicina, Instituto Universitario de Oncología, Universidad de Oviedo, Oviedo, Spain; Nuffield Department of Women’s and Reproductive Health, University of Oxford, Oxford, United Kingdom

## Abstract

Mitochondrial stress triggers a response in the cell’s mitochondria and nucleus, but how these stress responses are coordinated *in vivo* is poorly understood. Here, we characterize a family with myopathy caused by a dominant p.G58R mutation in the mitochondrial protein CHCHD10. To understand the disease etiology, we developed a novel knock-in mouse model and found that mutant CHCHD10 aggregates in affected tissues, applying a toxic protein stress to the inner mitochondrial membrane. Unexpectedly, survival of CHCHD10 knock-in mice depended on a protective stress response mediated by OMA1. The OMA1 stress response acted both locally within mitochondria, inhibiting mitochondrial fusion, and signaled outside the mitochondria, activating the integrated stress response. We additionally identified an isoform switch in the terminal complex of the electron transport chain as a novel component of this response. Our results demonstrate that OMA1 is essential for neonatal survival conditionally in the setting of inner mitochondrial membrane stress, coordinating local and global stress responses to reshape the mitochondrial network and proteome.

**Graphical Abtract:** 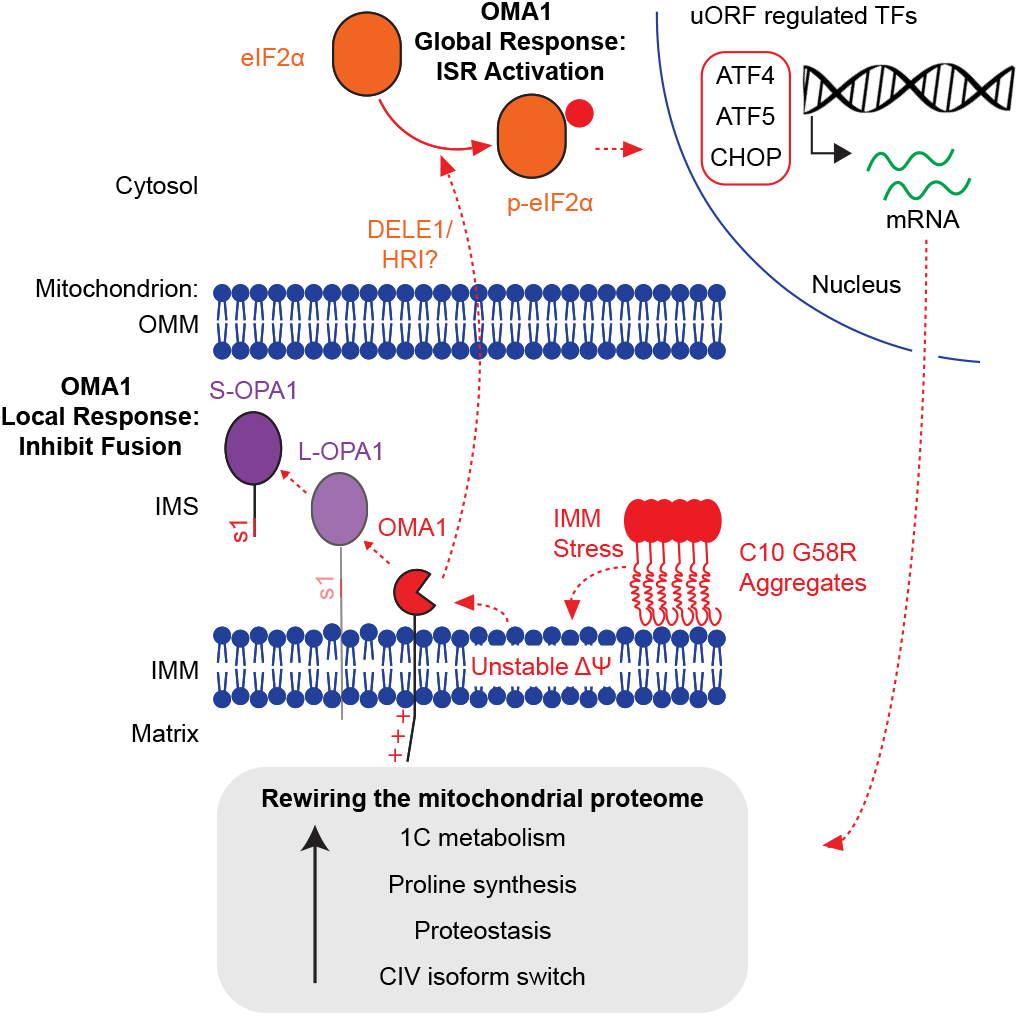

## Introduction

Mitochondria produce energy by coupling the transport of protons across the inner mitochondrial membrane (IMM) to the production of adenosine triphosphate (ATP) (1). The IMM additionally forms a barrier between the cytosol and the proinflammatory mitochondrial DNA (mtDNA) and RNA in the mitochondrial matrix (2–6). The integrity of the IMM is, thus, critical to mitochondrial function and tissue homeostasis.

IMM integrity is sensed by quality control pathways which include the IMM peptidase OMA1 and the PINK1/PARKIN pathway (7–10). OMA1 senses IMM integrity through its N-terminus, which responds to loss of IMM voltage by increasing OMA1 peptidase activity (11). Once activated by loss of IMM voltage, OMA1 cleaves itself and the IMM fusion protein L-OPA1. This leads to selective fragmentation and isolation of dysfunctional mitochondria. Thus, the OMA1-OPA1 axis acts as a local response to isolate dysfunctional mitochondria from the mitochondrial network.

Recently, OMA1 was additionally found to mediate a global response, communicating mitochondrial dysfunction to the cytosol and the nucleus through the integrated stress response (ISR) in cultured cells (12, 13). Along this axis, OMA1 cleaves the IMM protein DELE1, which retro-translocates to the cytosol to activate HRI, one of four eukaryotic translation initiation factor 2α (eIF2α) kinases. The eIF2α kinases are activated by diverse cellular stresses to phosphorylate eIF2α on S51, leading to a generalized stress response (14, 15). This response inhibits cytosolic translation of most mRNAs but increases translation of select mRNAs that contain upstream open reading frames, notably, the transcription factors ATF4, ATF5, and CHOP. These transcription factors upregulate pathways involved in one-carbon metabolism, amino acid synthesis, amino acid transport, and tRNA aminoacylation, among others, thereby restoring cellular homeostasis. Although mitochondrial stress leads to an ATF4 transcriptional response in diverse models of mitochondrial disease (16–22), it is not known whether OMA1 activation can mediate mitonuclear communication *in vivo*. Additionally, it is not known whether OMA1 activation may protect against specific mitochondrial stresses, as prior *in vivo* studies have found that OMA1 activation is often maladaptive (23, 24).

We recently discovered that OMA1 is activated in cell culture and *in vivo* in response to pathogenic mutations in the protein CHCHD10 (coiled-coil-helix-coiled-coil-helix domain containing 10; hereafter, C10) (20). C10 is a small mitochondrial intermembrane space (IMS) protein that extends into mitochondrial cristae and, together with its paralog CHCHD2 (hereafter, C2), is important for maintaining inner membrane integrity and electron transport chain (ETC) function (20, 25–29). Dominant mutations in C10 cause a spectrum of neuromuscular disorders that includes autosomal dominant isolated mitochondrial myopathy (IMMD; R15S/G58R) (30, 31), myopathy with amyotrophic lateral sclerosis/frontotemporal dementia (ALS/FTD; S59L) (32), adult-onset spinal muscular atrophy (SMA-J; G66V) (33), and ALS (R15L) (34), while the T61I mutation in C2 causes Parkinson’s disease (35). It is not resolved which C10 mutation, R15S or G58R, causes IMMD, as the two mutations co-segregated with disease in the one reported family (hereafter, the US family).

Most of these pathogenic mutations cluster within a highly conserved α-helix in the middle of the proteins and lead to C10 or C2 misfolding, suggesting a shared pathogenetic mechanism for these dominant mutations (16, 28, 36, 37). Additionally, knock-in (KI) mouse models of the C10 S59L mutation cause a more severe myopathy and cardiomyopathy than single C10 or C2/C10 double knockouts, suggesting that C10 mutant pathogenicity is due at least in part to toxic gain of function (16, 20, 27, 38). The basis of the selective tissue vulnerability to different C10 mutations, however, is not clear.

Here, we present a family with myopathy and cardiomyopathy due to the C10 G58R mutation (hereafter, the UK family), demonstrating that the G58R mutation is the cause of IMMD. This phenotype is also observed in a novel C10^G58R/+^ (hereafter, C10^G58R^) KI mouse model, corroborating that the G58R mutation is pathogenic. We further identify that OMA1 is critical for C10^G58R^ neonatal survival, orchestrating both local and global stress responses to IMM stress from C10 protein misfolding.

## Results

### The CHCHD10 G58R mutation causes autosomal dominant myopathy and cardiomyopathy

We identified a family with autosomal dominant mitochondrial myopathy and cardiomyopathy (Figure 1A). The mother of the UK family was previously described 50 years ago as having childhood-onset myopathy with mitochondrial inclusions of unknown cause (39). EMG showed only myopathic features with no fibrillation potentials at rest (39). The proband similarly had childhood onset myopathy with delayed motor milestones, inability to run or jump, positive Gowers’s sign, frequent falls, and lax ligaments (Figure 1B). By the age of 20 years the proband was wheelchair-bound and had received a heart transplant due to severe progression of his cardiomyopathy. He died from lymphoma soon afterwards, likely as a complication of immunosuppression. The proband had an unaffected sister (III-1) and a brother with myopathy and cardiomyopathy who died in childhood (III-2).

**Figure 1.**
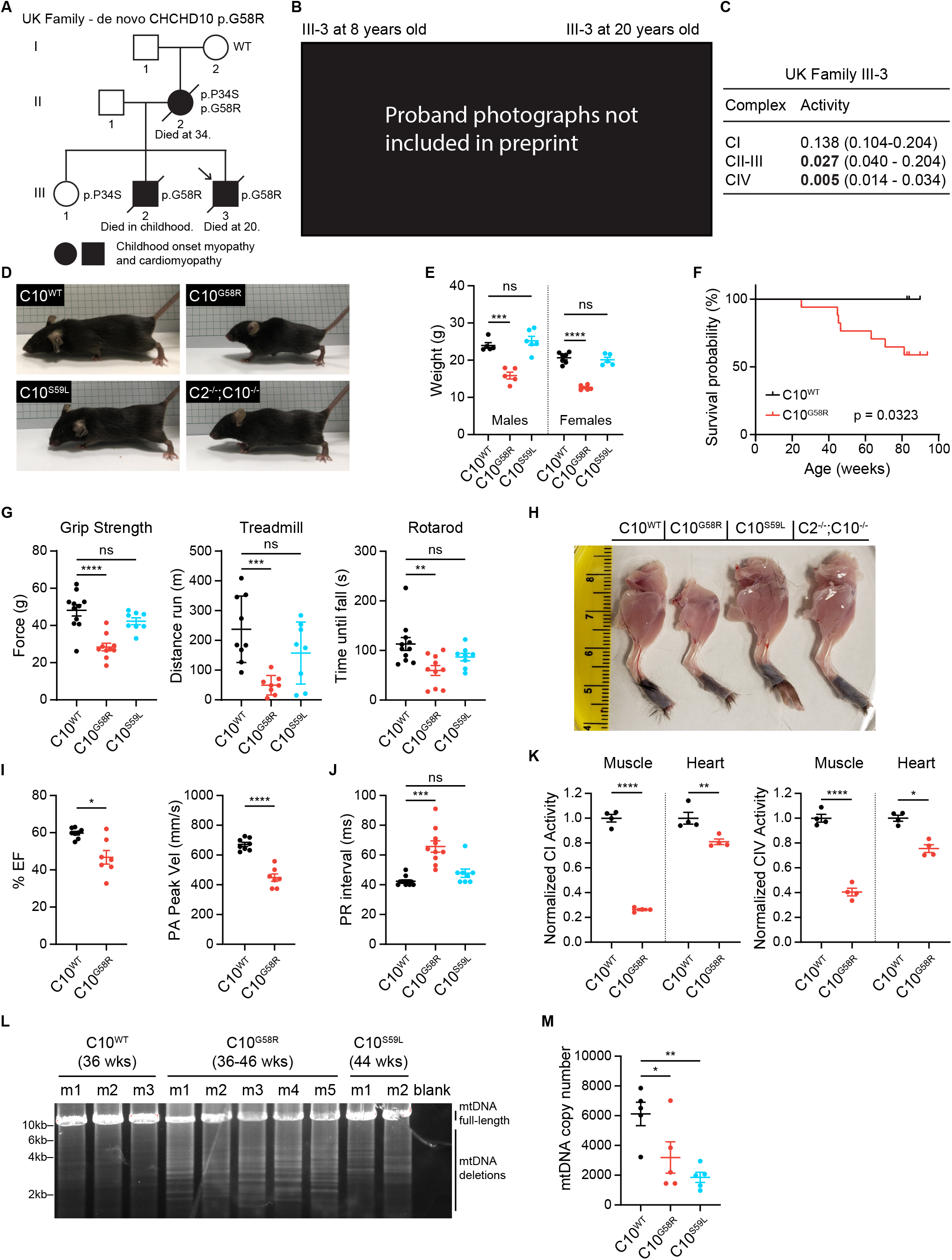
The C10 G58R mutation causes dominant myopathy and cardiomyopathy in human and mouse. (A) Pedigree of the UK family. Arrow indicates the proband. (B) The proband (III-3) at 8 (left) and 20 (right) years old. (C) ETC complex activities from the proband’s pectoralis (III-3). (D) 15-week-old C10^WT^ and C10^G58R^ mouse littermates, with C10^S59L^ and C2/C10 DKO mice of the same age for comparison. (E) Weights of 10–11-week-old C10^WT^, C10^G58R^, and C10^S59L^ mice. (F) Survival curve of C10^WT^ (n = 9) and C10^G58R^ (n = 17) mice. (G) Grip strength, treadmill fatigue test, and rotarod assays of 18-week-old C10^WT^ and C10^G58R^ mice, and 25-week-old C10^S59L^ mice. (H) Skinned hindlimbs of 15-week-old C10^WT^, C10^G58R^, C10^S59L^, and C2/C10 DKO mice. (I) Percent ejection fraction (EF) and pulmonary artery (PA) peak velocity from C10^WT^ and C10^G58R^ mouse echocardiography. (J) PR intervals from C10^WT^, C10^G58R^, and C10^S59L^ mouse electrocardiograms. (K) Complex I (CI) and IV (CIV) activities from C10^WT^ and C10^G58R^ mouse heart and tibialis lysates. (L) Long-range PCR of a 12.8kb segment of C10^WT^, C10^G58R^, and C10^S59L^ mouse heart mtDNA. Abbreviation – m: mouse. (M) mtDNA copy number of 36–46-week-old C10^WT^, C10^G58R^, and C10^S59L^ mouse hearts. Each datapoint represents a mouse. Error bars represent SEM. *p<0.05, **p<0.01, ***p<0.001, ****p<0.0001. See also Figures S1 and S2, and Supplemental File 1.

Whole exome sequencing demonstrated a p.G58R mutation in the proband, and Sanger sequencing of C10 in other members of the UK family demonstrated the p.G58R variant *in trans* with the benign p.P34S variant in the mother, the benign p.P34S variant in the unaffected daughter, and the p.G58R variant in the affected brother. The maternal grandmother had neither the p.G58R nor the p.P34S variant, suggesting that the mother most likely inherited the p.P34S variant from the maternal grandfather and the p.G58R variant most likely arose in the mother *de novo*. Together, these findings establish that the G58R variant is pathogenic and suggest that it primarily affects striated muscle in humans.

Assessment of muscle tissue from members of the UK family was also consistent with mitochondrial myopathy and similar to findings reported for the US family (30, 31). Biopsy of the proband’s pectoralis muscle at age 8 showed excess lipid droplets in type I fibers, suggestive of a mitochondrial myopathy (Figure S1A). mtDNA deletions were present at a low level in the pectoralis muscle but not the heart (detectable by long-range PCR but not Southern blot) (Figure S1B). Mitochondrial complex activity assays of his muscle showed normal complex I activity, moderately decreased complex II-III activity, and more severely decreased complex IV activity (Figure 1C). Additionally, the mother’s muscle biopsy showed a predominance of type I fibers, with ragged-red fibers and cytochrome c oxidase (COX)-negative fibers (Figure S1C).

To further study the pathogenesis of C10 G58R, we generated a KI mouse model harboring the homologous mutation in the corresponding mouse gene (p.G54R in mouse, but the human numbering will be used throughout) (Figure S1D). C10^G58R^ mice were smaller, had decreased body weight, and died prematurely (Figures 1D - F). The weight difference for C10^G58R^ was already clear at the time of weaning (Figure S1E). By contrast, C10^S59L/+^ (hereafter, C10^S59L^) and C2/C10 double knockout (DKO) mice were similar in size to WT at 15 weeks (Figure 1D). Total food intake did not differ between C10^G58R^ and C10^WT^ mice, and there was no difference in the percent of lean and fat mass between the genotypes (Figures S2A and B). The respiratory exchange ratio, measured by Comprehensive Laboratory Animal Monitoring System (CLAMS), tended to be higher for C10^G58R^ mice at night (indicating increased carbohydrate reliance) and lower during the day (indicating increased fatty acid reliance) compared to C10^WT^ littermates, although these differences did not reach statistical significance (Figure S2C). When accounting for body weight, there was no difference between C10^WT^ and C10^G58R^ mice in oxygen consumption, carbon dioxide production, or energy expenditure (Figure S2D).

Similar to the proband, C10^G58R^ mice had myopathy, with decreased grip strength, worse rotarod performance, and increased treadmill-induced fatigue when tested at 18 weeks (Figure 1G). C10^S59L^ mice tested at a slightly older age (25 weeks) did not yet differ from C10^WT^ (Figure 1G), although both C10^G58R^ and C10^S59L^ mice took longer to descend a 50 cm pole (Figure S1F). Consistent with the poor motor function, C10^G58R^ mice had smaller leg muscles compared to C10^WT^, C10^S59L^, and C2/C10 DKO mice (Figure 1H), and muscle fiber size was significantly smaller in C10^G58R^ mice than in C10^WT^ mice (Figures S1G and H), but no COX-negative fibers were observed (Figure S1G). Additionally, C10^G58R^ (but not C10^S59L^) mouse tibialis muscles had increased lipid droplets, phenocopying muscle biopsy findings from members of the UK family (Figures S1G and I). Also similar to the proband, C10^G58R^ mice had decreased heart function measured by echocardiography (Figure 1I and Supplemental File 1), and additionally had atrioventricular heart block (Figures 1J and S1J). Consistent with a mitochondrial basis for heart and muscle dysfunction, complex I and complex IV activities were diminished in C10^G58R^ mouse heart and muscle (Figure 1K), and C10^G58R^ mice had multiple mtDNA deletions and decreased mtDNA copy number in the heart, consistent with a mild defect in mtDNA maintenance (Figures 1L and M). Together, these results demonstrate that the C10^G58R^ mouse model recapitulates the myopathy and cardiomyopathy phenotypes of the UK family and has a more severe myopathy than the C10^S59L^ mouse model.

### CHCHD10 G58R and S59L mutations form distinct aggregates

We next explored the basis for phenotypic differences between C10^G58R^ and C10^S59L^ mice. Analyzing C10’s amino acid sequence with EVcouplings and its predicted structure with EVfold (40, 41), we found that the G58 and S59 residues are highly conserved and are predicted to introduce a break in the central α-helix (Figures 2A and B). Notably, while the S59L mutation increased the calculated hydrophobicity of the region, as recognized previously (16), the G58R mutation decreased it, suggesting that mutations in these neighboring residues may have differential physiochemical effects on the α-helix (Figure 2B).

**Figure 2.**
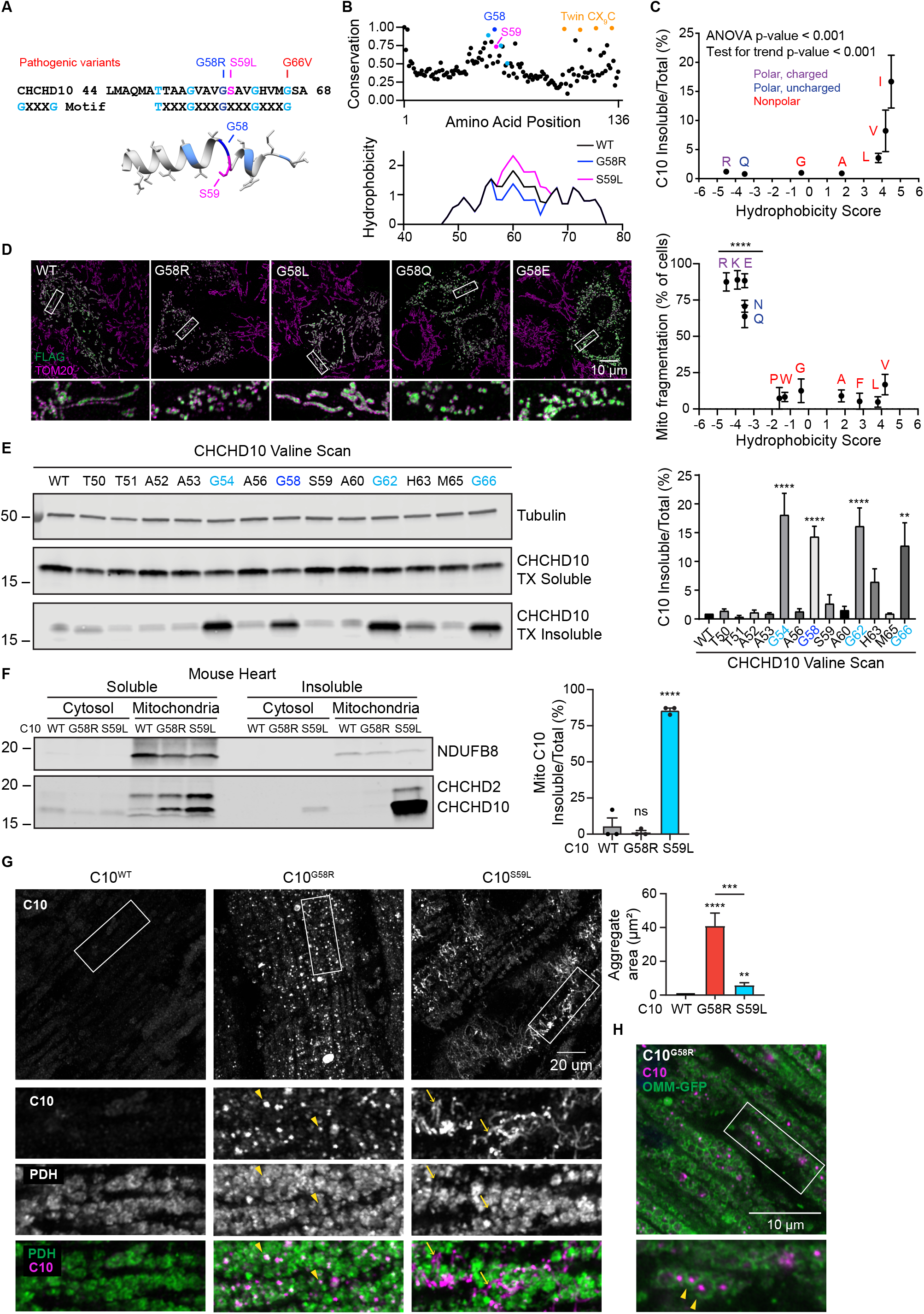
C10 G58R is predominantly soluble but forms aggregates that are distinct from C10 S59L. (A) Top: amino acid sequence of the hydrophobic α-helix of C10 showing pathogenic variants and highlighting the GXXXG motif. Bottom: EVfold prediction of the structure of the α-helix. (B) Top: EVcouplings conservation analysis of C10. Bottom: predicted hydrophobicity of the region around the α-helix. (C) Levels of insoluble/total C10 in HEK293 C2/C10 DKO cells after transfection with C10 containing G58 substitutions with amino acids of varying hydrophobicity on the Kyte-Doolittle scale (x-axis), (n = 3 biological replicates). (D) Left: representative Airyscan images of mitochondria of HeLa cells transfected with C10 containing the indicated G58 substitutions. Right: quantification of mitochondrial fragmentation in HeLa cells transfected with C10 containing G58 substitutions with amino acids of varying hydrophobicity, (n ≥ 50 cells per replicate from 3 biological replicates). (E) Left: representative blot of a Triton-X (TX)-soluble/insoluble assay of HEK293 C2/C10 DKO cells transfected with C10 whereby individual residues of the α-helix were mutated to valines. Right: quantification of the blots (n = 3 biological replicates). (F) Left: representative immunoblot of a soluble/insoluble assay of C10 from the cytosolic or mitochondrial fraction of C10^WT^, C10^G58R^, and C10^S59L^ mouse hearts. Right: quantification of the blots, (n = 3 mice per genotype). (G) Left: Airyscan images of 15-week-old C10^WT^, C10^G58R^, and C10^S59L^ mouse hearts stained for C10 and PDH. Arrowheads show intramitochondrial aggregates and arrows show extramitochondrial aggregates. Right: quantification of C10 aggregate area per field of view (FOV) of 36–46-week-old mice, (n = 3 mice C10^WT^ and C10^S59L^, n = 4 mice C10^G58R^; 8 FOV per mouse). (H) Representative Airyscan image of a 12-week-old C10^G58R^; mito-QC^tg/+^ mouse heart showing intramitochondrial C10 aggregates. Abbreviation – OMM: outer mitochondrial membrane, (n = 2 mice). Error bars represent SEM. *p<0.05, **p<0.01, ***p<0.001, ****p<0.0001. See also Figure S3.

Consistent with its conservation, we found that the G58 residue was intolerant to substitution when exogenously expressed in cultured cells. Hydrophobic substitutions decreased C10 protein solubility (Figures 2C and S3A), whereas large hydrophilic substitutions caused mitochondrial fragmentation, similar to what we and others previously reported for the pathogenic G58R substitution (Figure 2D) (20, 28, 30). An arginine scan of this region showed that the G58 position is the most prone to causing mitochondrial fragmentation (Figure S3B). We previously observed that the neuronopathy-causing G66V mutation also causes C10 insolubility (28). As the G58 and G66 residues lie on either end of a GXXXGXXXG motif, we systematically assessed the effect of valine substitutions on C10 solubility. Notably, substitution of any glycine residue strongly reduced C10 solubility (Figure 2E). Thus, increasing hydrophobicity of the glycine face of the α-helix may promote an insoluble C10 conformation. This contrasts with the C10 G58R substitution, which may adopt a soluble conformation that is more prone to inducing mitochondrial fragmentation.

The S59L mutation was previously reported to form insoluble C10 aggregates in mouse heart (16, 20), so we compared the solubility and aggregation tendency of the G58R and S59L mutations in this tissue. Consistent with the data from cultured cells and the calculated change in hydrophobicity, C10 with the G58R mutation was soluble, in contrast to C10 with the S59L mutation (Figure 2F). C10 levels were much higher in C10^S59L^ than in C10^G58R^ mouse hearts, raising the possibility that insolubility may drive the increased protein level of C10 S59L. C10 protein aggregates were prominent in C10^G58R^ mouse hearts, but had a distinct morphology and localization compared to those in C10^S59L^ mouse hearts (Figure 2G). While C10 S59L protein tended to form extramitochondrial, filamentous aggregates, C10 G58R protein formed intramitochondrial, punctate aggregates (Figures 2G and H). Additionally, C10 G58R aggregates occupied a larger area of the tissue in heart and in skeletal muscle (Figures 2G and S3C). Immuno-EM of C10 G58R protein expressed in HEK293 C2/C10 DKO cells showed that it localized to the IMS side of cristae, similar to endogenous C10 WT protein (Figure 3A) (28). Together, these findings suggest that C10 G58R and S59L proteins likely form distinct aggregates, which may underlie their differential toxicities.

**Figure 3.**
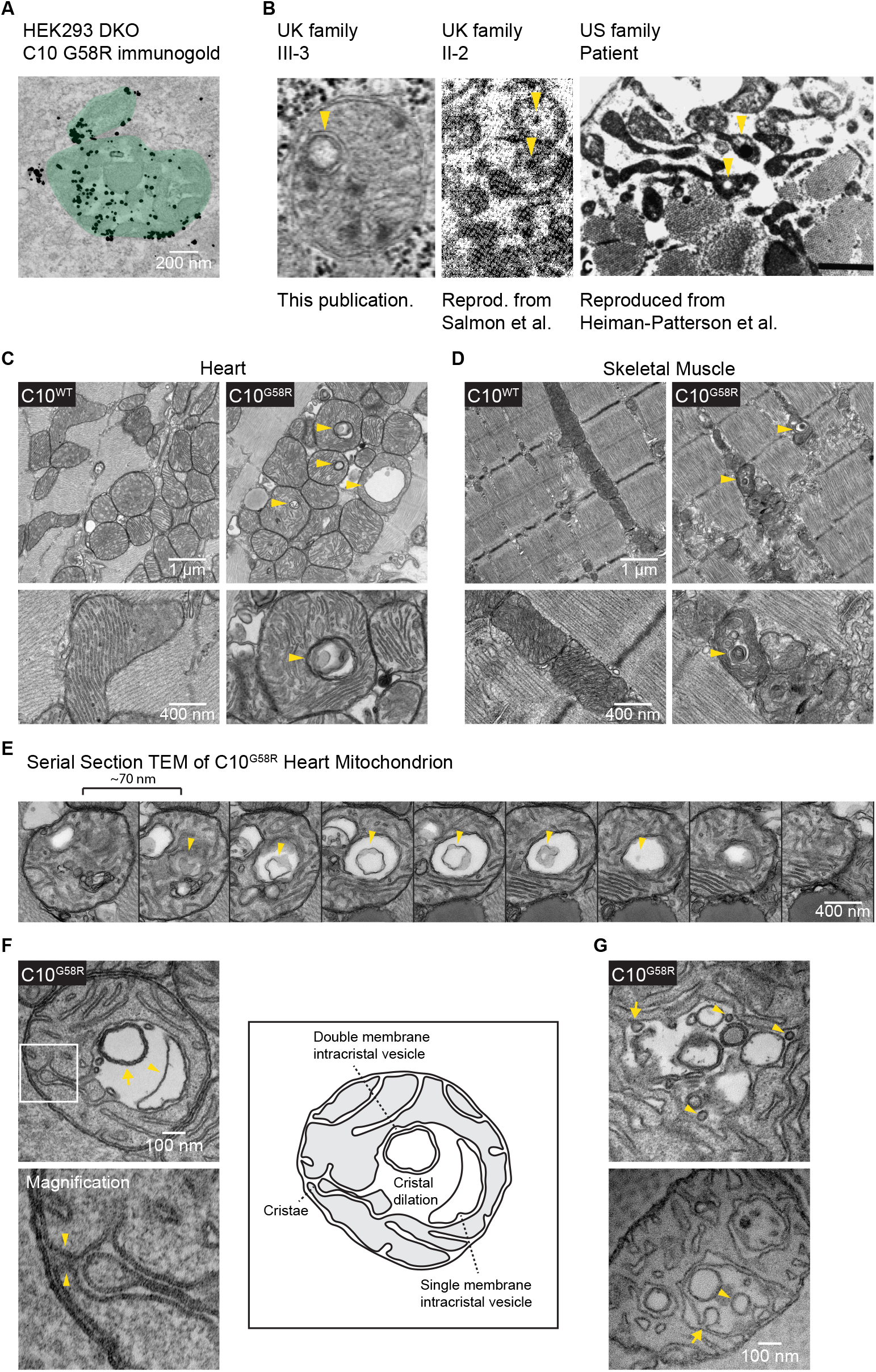
Intracristal inclusions are characteristic of affected C10 G58R patient and mouse muscle. (A) Immunogold-EM of C10 G58R overexpressing (OE) C2/C10 DKO HEK293 cells, (representative of ≥ 5 mitochondria in 1 biological replicate). (B) Intramitochondrial inclusions in muscles of patients with the G58R mutation. (C) TEM of 28-week-old C10^WT^ and C10^G58R^ mouse hearts. Arrowheads indicate intramitochondrial inclusions, (representative of ≥ 15 fields in 2 biological replicates). (D) TEM of tibialis from 14-week-old C10^WT^ and C10^G58R^ mice on the OMA1^+/-^ background. Arrowheads indicate intramitochondrial inclusions, (representative of ≥ 10 fields in 1 biological replicate). (E) Serial TEM following an intramitochondrial inclusion containing a membranous vesicle within, (representative of ≥ 6 mitochondria from 1 biological replicate). (F) Top: TEM of an ultrathin section of a 33-week-old C10^G58R^ mouse heart showing cristal dilation, a single-membraned intracristal vesicle (arrowhead), and a double-membraned intracristal vesicle (arrow). Bottom: magnification showing continuity with the IMM indicated by arrowheads. Right: sketch and labels of the image on the left, (representative of ≥ 10 mitochondria in 1 biological replicate). (G) TEM of an ultrathin section of a 33-week-old C10^G58R^ mouse heart showing inner membrane active budding events (arrows) and completed budding (arrowheads), (representative of ≥ 5 mitochondria in 1 biological replicate).

### Intracristal inclusions are characteristic of C10 G58R pathology and likely reflect inner membrane stress

Double-membraned inclusions within mitochondria were observed in the proband’s muscle biopsy and were similar in appearance to the “inclusion bodies” previously noted in his mother and “globular inclusions” previously noted in a member of the US family (Figure 3B) (31, 39). In C10^G58R^ mouse heart and skeletal muscle (but not C10^WT^ or C10^S59L^ tissues), similar inclusions were observed in nearly every field of view (Figures 3C and D). These inclusions were enclosed within mitochondria, as demonstrated by serial EM (Figure 3E), and were formed from the cristal membrane with which they were continuous (Figure 3F). Thus, they represented dilations of cristae, which sometimes contained membranous intracristal vesicles with single or multiple membranes (Figure 3F). Additionally, we observed possible inward budding events of cristal membrane into the intracristal space, suggesting that at least some of the internal membranous structures may be derived from the cristal membrane (Figure 3G). Together, these observations establish that intracristal inclusions are characteristic of C10 G58R myopathy.

### OMA1 is activated by IMM stress in C10^G58R^ mice and is essential for their survival

We reasoned that the cristal inclusions in C10^G58R^ striated muscle may be symptomatic of inner membrane stress exerted by C10 G58R protein misfolding in the IMS. We next assessed whether OMA1, a peptidase monitoring IMM stress, might be activated by the G58R mutation *in vivo*, similar to what we reported previously for the G58R mutation in cell culture and the S59L mutation *in vivo* (20).

OMA1 cleavage of OPA1 was increased in C10^G58R^ mice in all tissues examined, as reflected by a decrease in OPA1 long forms (L-OPA1, bands a and b) and an increase in OMA1-generated short forms (S-OPA1, bands c and e) (Figures 4A and B, and S4A). OMA1 was more strongly activated by C10 G58R than by C10 S59L in all tissues, despite higher protein levels of C10 in C10^S59L^ mouse tissue (Figures 4A and C). As expected, OMA1 levels were negatively correlated with OPA1 processing, due to OMA1 autocatalysis upon activation (Figure 4A) (11). OMA1 cleavage of L-OPA1 was also observed in the heart and skeletal muscle tissue of the proband, demonstrating that the OMA1 stress response to C10 G58R protein is conserved between mice and humans (Figure 4D). These results establish that the C10 G58R mutation (and to a lesser extent the S59L mutation) cause widespread OMA1 activation *in vivo*.

**Figure 4.**
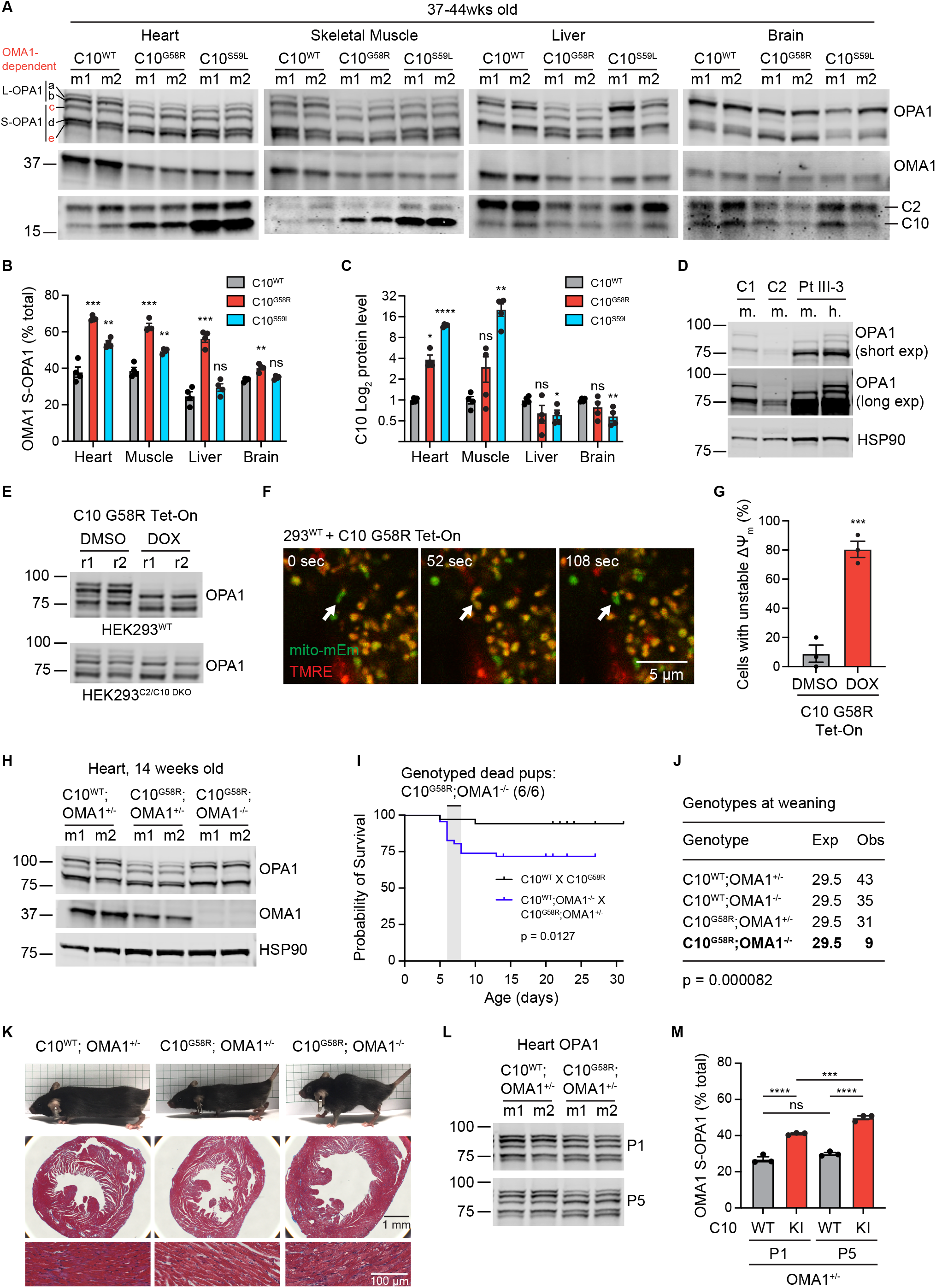
OMA1 is activated in C10 G58R mouse and patient tissues and is necessary for survival. (A) Immunoblots of OPA1, OMA1, C2, and C10 from different tissues of 37–44-week-old C10^WT^, C10^G58R^, and C10^S59L^ mice. Loading controls in Figure S4A. (B) Quantification of OMA-cleaved OPA1 bands (c+e/total) from (A), (n = 4 mice per genotype). (C) Quantification of C10 levels from (A). (D) Immunoblot of OPA1 levels from the C10 G58R proband (patient III-3) and two non-myopathic controls. Abbreviations – C: control, m: muscle, h: heart. (E) Immunoblot of OPA1 levels from Tet-On HEK 293 cells treated overnight with DMSO or DOX (overexpressing C10 G58R) on either a WT background (top) or C2/C10 DKO background (bottom). Abbreviation – r: replicate, (n ≥ 3 biological replicates). (F) Representative confocal time series of Tet-On HEK293 cells overexpressing C10 G58R. Arrow indicates a mitochondrial flicker event. (G) Quantification of the cells from (F) containing mitochondria with unstable ΔΨ_m_ after overnight treatment with DMSO or doxycycline, (n ≥ 10 mitochondria per replicate in 3 biological replicates per condition). (H) Immunoblot of OPA1 and OMA1 levels from 14-week-old littermates of the C10^WT^ ; OMA1^-/-^ X C10^G58R^ ; OMA1^+/-^ cross. Abbreviation – m: mouse, (representative of n = 4 mice per genotype). (I) Survival curves of pups from the C10^WT^ X C10^G58R^ and C10^WT^ ; OMA1^-/-^ X C10^G58R^ ; OMA1^+/-^ crosses. All 6 dead pups genotyped from the latter cross were C10^G58R^ ; OMA1^-/-^. (n = 18 pups from C10^WT^ X C10^G58R^ and n = 46 pups from C10^WT^ ; OMA1^-/-^ X C10^G58R^ ; OMA1^+/-^). (J) Expected vs observed numbers of pup genotypes from the C10^WT^ ; OMA1^-/-^ X C10^G58R^ ; OMA1^+/-^ cross. (K) Top: 14-week-old littermates from the C10^WT^ ; OMA1^-/-^ X C10^G58R^ ; OMA1^+/-^ cross. Middle: Masson’s trichrome (MT) OMA1 protects against inner mitochondrial membrane stress stain of mouse hearts. Bottom: magnified MT stain of heart showing prominent vacuolation in C10^G58R^ ; OMA1^-/-^, (representative of n = 3 mice per genotype). (L) Immunoblot of OPA1 levels from hearts of C10^WT^ or C10^G58R^ mice on the OMA1^+/-^ background, on postnatal days 1 and 5, (representative of n = 3 mice per genotype). (M) Quantification of OMA-cleaved OPA1 bands (c+e/total) from (L). Error bars represent SEM. *p<0.05, **p<0.01, ***p<0.001, ****p<0.0001. See also Figure S4.

We next examined the mechanism of OMA1 activation by the C10 G58R mutation in HEK293 cells. We found that overnight C10 G58R expression activates OMA1 on a WT background, as we reported previously (20), as well as on a C2/C10 DKO background, demonstrating that OMA1 activation by C10 G58R is at least in part due to a toxic gain of function (Figure 4E). OMA1 can be activated by sustained loss of the voltage potential across the IMM (ΔΨ_m_) or by ΔΨ_m_ instability (also known as mitochondrial flicker) (42, 43). Although acute C10 G58R expression did not decrease total cell ΔΨ_m_ as measured by flow cytometry (Figure S4B), it caused ΔΨ_m_ instability and mitochondrial fragmentation observed by live cell imaging (Figures 4F and G, S4C, and Supplemental Video 1), suggesting that the G58R mutation activates OMA1 by causing ΔΨ_m_ instability.

Having established that OMA1 is activated by the G58R mutation, we next asked if the OMA1 stress response is protective or maladaptive. We crossed C10^G58R^ mice to OMA1 KO mice, which have a normal lifespan and tend toward increased weight (Figure 4H) (44). To our surprise, the G58R mutation was neonatal lethal on the OMA1 KO background, with few C10^G58R^ ; OMA1^-/-^ mice surviving to weaning (Figures 4I and J, and S4D). Most of the C10^G58R^ ; OMA1^-/-^ pups died between postnatal days (P) 5-8, with the few escapees dying by 15 weeks of age. Notably, the escapees had markedly enlarged hearts despite decreased body size, in contrast to the double heterozygous C10^G58R^; OMA1^+/-^ mice, which had decreased heart size in proportion with their body size (Figure 4K). OMA1 KO increased the number of p62 aggregates but not C10 aggregates in C10^G58R^ mouse hearts (Figure S4E) and worsened the mtDNA deletion load (Figure S4F).

To further investigate the neonatal lethality of this cross, we examined timed pregnancies. C10^G58R^ ; OMA1^-/-^ mice were grossly indistinguishable from their littermates at embryonic day 18.5 and P5 (Figures S4G and H) and did not differ by body weight at P1 (Figure S4I). However, their body weight dropped at P5 (Figure S4I), indicating decompensation. In the heart, OMA1 was already activated at P1 and OMA1 activity increased at P5 (Figures 4L and M). In marked contrast to the C10^G58R^; OMA1^+/-^ X OMA1^-/-^ cross, the C10^S59L^; OMA1^+/-^ X OMA1^-/-^ cross was not neonatal lethal (Figure S4J), suggesting that the OMA1 stress response is more critical for G58R survival. Together these results demonstrate that the OMA1 stress response is essential for G58R survival, with decompensation starting around P5.

### OMA1 fragments heart mitochondria in response to C10 G58R stress

OMA1 cleavage of L-OPA1 inhibits mitochondrial fusion, causing mitochondrial fragmentation (7, 8, 45). We previously observed that exogenously expressed C10 G58R fragments mitochondria in HeLa cells by activating OMA1 (20). Consistently, we observed that mitochondrial fragmentation from C10 mutants identified in Figures 2D and S3B also depended on OMA1 (Figures 5A and B). We additionally noted that exogenously expressed C10 G58R caused narrowing of the mitochondrial caliber in OMA1 KO cells (Figures 5A and C), similar to what had been reported in DRP1 deficient OMA1 KO cells (42). This may reflect matrix collapse due to ΔΨ_m_ instability across the IMM, as suggested for DRP1 deficient cells (42).

**Figure 5.**
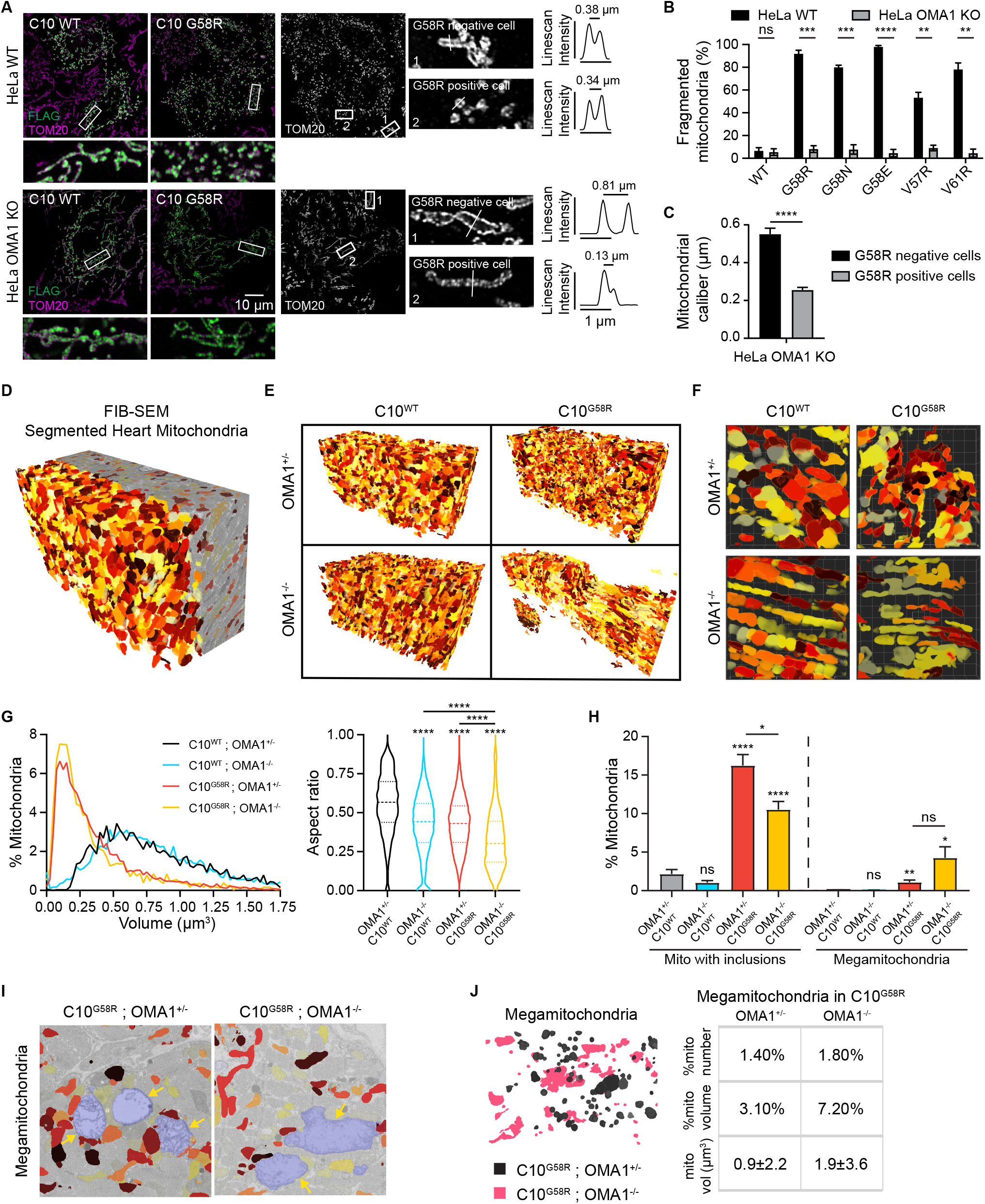
The C10 G58R mutation fragments heart mitochondria in an OMA1-dependent manner. (A) HeLa cells transfected with C10 WT and C10 G58R constructs on the WT (top) or OMA1 KO (bottom) background. Magnifications depict mitochondrial caliber, with TOM20 line scan intensities on the right. Images were taken with a Zeiss Airyscan microscope. (B) Mitochondrial fragmentation levels in HeLa cells transfected with C10 mutants on the WT or OMA1 KO background, (n ≥ 50 mitochondria per replicate in 3 biological replicates per condition). (C) Quantification of mitochondrial caliber in (A), (n = 42 mitochondria from 3 biological replicates). (D) Reconstruction of segmented FIB-SEM mouse heart mitochondria. Colors are arbitrary and only to aid visualization. (E) Complete datasets of 3D-reconstructed heart mitochondria of 14-week-old mice of the indicated genotypes. (F) Magnification of representative fields from the datasets in (E). (G) Volume distribution and aspect ratio measurements of mitochondria from (E), (n > 3159 mitochondria for each genotype from 1 biological replicate). (H) Percent of mitochondria with inclusions and percent of megamitochondria from TEM images of 14-week-old mouse hearts, (n = 3 mice per genotype for all genotypes but OMA1^+/-^ ; C10^G58R^, for which n = 2. 15 FOV quantified per mouse, with >1900 mitochondria assessed per genotype). (I) Megamitochondria (shaded in blue) from the FIB-SEM stack. Non-megamitochondria are colored for comparison. (J) Left: overlay of segmented megamitochondria from C10^G58R^ mice on the OMA1^+/-^ (black) or OMA1^-/-^ (pink) background. Right: percentage of megamitochondria in the FIB-SEM stack, percentage of megamitochondrial volume from total mitochondrial volume, and average volume of individual megamitochondria, (n = 119 OMA1^+/-^ ; C10^G58R^ megamitochondria, n = 58 OMA1^-/-^ ; C10^G58R^ megamitochondria). Error bars represent SEM. For the violin plots, the 25^th^ quartile, median, and 75^th^ quartile are indicated. *p<0.05, **p<0.01, ***p<0.001, ****p<0.0001. See also Figure S5.

To assess the effects of G58R and OMA1 activation *in vivo*, we analyzed heart mitochondria in a surviving 14-week-old C10^G58R^ ; OMA1^-/-^ mouse and its littermates by focused ion beam - scanning electron microscopy (FIB-SEM), followed by segmentation and 3D reconstruction of heart mitochondria (Figure 5D, Supplemental Video 2). C10^G58R^ ; OMA1^+/-^ mitochondria were smaller than C10^WT^ ; OMA1^+/-^ mitochondria (Figures 5E, F, and G), reflecting mitochondrial fragmentation. C10^WT^ ; OMA1^-/-^ mitochondria had the same average volume as C10^WT^ ; OMA1^+/-^, but they tended to be longer, as reflected by the decreased aspect ratio (Figure 5G). Interestingly, C10^G58R^ ; OMA1^-/-^ mitochondria were also longer but tended to have decreased caliber and thus smaller volume, similar to what was seen in cell culture with exogenous C10 G58R expression. They also had the lowest aspect ratio, reflecting their long, thin morphology. These data demonstrate that OMA1 activation causes mitochondrial fragmentation *in vivo*.

Next, we analyzed heart tissue from these genotypes by transmission electron microscopy (TEM), which has superior lateral resolution. As expected, there were more inclusions in the C10^G58R^ mutants on either background compared to C10^WT^ mouse hearts (Figure 5H). Notably, OMA1 KO partially blocked inclusion formation, suggesting that cleavage of L-OPA1 from the inner membrane may facilitate (but is not required for) their formation (Figure 5H).

Additionally, in C10^G58R^ mutants on both backgrounds, we observed megamitochondria: massively swollen mitochondria with lighter matrix density compared to surrounding mitochondria (Figures 5H and S5A), similar to what has been observed in other models of mitochondrial or metabolic stress (46). These megamitochondria were also seen in the FIB-SEM datasets of the C10^G58R^ mutants (Figure 5I and Supplemental Video 3). While the proportion of megamitochondria was similar for the two genotypes, the overall morphology of the megamitochondria (e.g., long or round) reflected that of the non-megamitochondria from their respective genotypes (Figure 5J). Taken together, these findings demonstrate that OMA1 fragments mitochondria in response to the G58R mutation *in vivo* and permits reshaping of the inner membrane to form inclusions.

### OMA1 signals mitochondrial stress through the ISR

A stress response involving ATF4 has previously been observed in both C10^S59L^ and C2/C10 DKO mice (as well as other mitochondrial myopathy models), although the mechanism for this response *in vivo* has not been clear (16, 20). ATF4 is often activated downstream of eIF2α phosphorylation in what is known as the ISR. To see if the ISR was activated in our C10^G58R^ and C10^S59L^ models, we immunoblotted for eIF2α phosphorylation. Levels of p-eIF2α / eIF2α were significantly increased in C10^G58R^ and C10^S59L^ hearts and C10^G58R^ muscle compared to C10^WT^, demonstrating ISR activation (Figures 6A and B).

**Figure 6.**
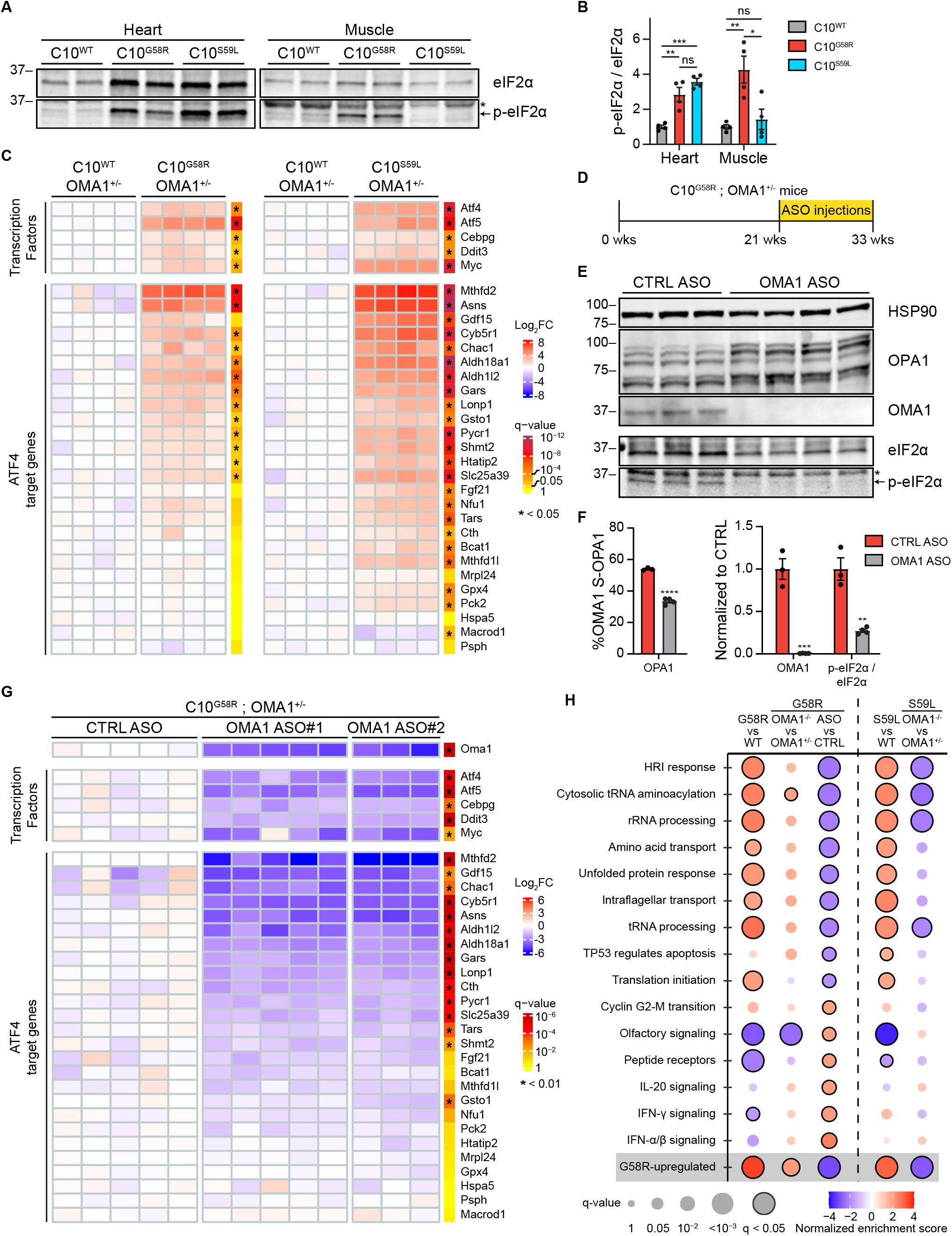
OMA1 signals mitochondrial stress to activate the integrated stress response. (A) Representative immunoblot of eIF2α and p-eIF2α from heart and muscle lysates of 37–44-week-old C10^WT^, C10^G58R^, and C10^S59L^ mice. *Nonspecific band. (B) Quantification of (A), (n = 4 mice per genotype). (C) Microarray data of hearts from 14-week-old C10^G58R^ vs C10^WT^ and 20-week-old C10^S59L^ vs C10^WT^ mice, all on the OMA1^+/-^ background. Each column represents a mouse, and q-values represent genome-wide significance. (D) Timeline of the ASO experiment. (E) Immunoblots of OPA1, OMA1, eIF2α, and p-eIF2α of C10^G58R^ mouse hearts on the OMA1^+/-^ background treated with either a nontargeting (CTRL) or OMA1 ASO. Loading controls for eIF2α and p-eIF2α in Figure S6F. *Nonspecific band. (F) Quantification of (E), (n = 3 mice CTRL ASO and n = 4 mice OMA1 ASO). (G) Microarray data of 33-week-old C10^G58R^ mouse hearts on the OMA1^+/-^ background treated with either a nontargeting (CTRL) or OMA1 ASO. Each column represents a mouse, and q-values represent genome-wide significance. (H) GSEA of reactome pathways of the listed comparisons. Pathways shown are all the pathways with a q-value < 0.025 in the C10^G58R^ OMA1 ASO vs CTRL ASO comparison. Error bars represent SEM. *p<0.05, **p<0.01, ***p<0.001, ****p<0.0001. See also Figure S6 and Supplemental Files 2 and 3.

To examine the ISR further, we looked at global gene expression in C10^G58R^ and C10^S59L^ mouse hearts on the OMA1^+/-^ background (Supplemental File 2). Gene set enrichment analysis (GSEA) of reactome pathways showed that EIF2AK1 (HRI) response, tRNA aminoacylation, and mitochondrial translation were among the most upregulated pathways in C10^G58R^ vs C10^WT^ hearts (Supplemental File 3). Indeed, we found that transcription factors involved in the ISR (ATF4, ATF5, CHOP, and CEBPG) were all significantly upregulated in both C10^G58R^ and C10^S59L^ mouse hearts (Figure 6C). Furthermore, many ATF4 target genes – including MTHFD2, ASNS, LONP1, and ALDH18A1 – were found to be upregulated in both C10 mutant strains. This confirmed that the ISR is activated in both C10^G58R^ and C10^S59L^ mouse hearts.

We next asked if OMA1 mediates the ISR *in vivo*, similar to what was recently reported in cultured cells (12, 13). In addition to analyzing the surviving C10^G58R^ ; OMA1^-/-^ mice, which were decompensated as discussed above, we knocked down OMA1 in adult C10^G58R^ ; OMA1^+/-^ mice using either nontargeting or one of two OMA1-specific antisense oligomers (ASOs) (Figure 6D). In contrast to the constitutive OMA1 KO, knockdown of OMA1 in the adult animals was relatively well-tolerated over the 12-week treatment, with only two out of nine OMA1 ASO-treated animals dying, and no or mild effects on cardiac function, motor function, and mtDNA stability (Figures S6A-E and Supplemental File 1). Immunoblotting confirmed knockdown of OMA1 and partial restoration of the long (noncleaved) forms of OPA1 in ASO-treated mouse hearts (Figures 6E and F). As the two OMA1 ASOs performed similarly, they were grouped together for analysis.

Levels of p-eIF2α / eIF2α were decreased to 27% of control ASO following OMA1 ASO treatment (Figures 6E and F, and S6F), indicating inhibition of the ISR. Similarly, ISR transcription factors and downstream ATF4-regulated genes were reduced in the OMA1 ASO vs. control ASO comparison (Figure 6G). Consistently, EIF2AK1 (HRI) response to heme deficiency emerged as the most downregulated reactome pathway by GSEA (Figure 6H and Supplemental File 3). Altogether, ∼70% of the differentially expressed genes (DEGs) identified in the C10^G58R^ vs C10^WT^ comparison were significantly suppressed by OMA1 ASO, suggesting that OMA1 drives most of the transcriptional response to the G58R mutation (Figure S6G). Similar to OMA1 knockdown in the C10^G58R^ mice, constitutive KO of OMA1 in C10^S59L^ mice suppressed the ISR (Figures S6H and I). By contrast, constitutive KO of OMA1 in C10^G58R^ mice mildly increased expression of some ATF4-dependent genes, likely reflecting alternative activation of ATF4 in the decompensated mice (Figures S6H and I). These data demonstrate that OMA1 signals IMM stress through the ISR *in vivo*.

### OMA1 activation shapes the mito-proteome through mitonuclear signaling

We next examined how the G58R mutation affects the heart mitochondrial proteome by label-free quantification mass spectrometry. OMA1 levels were decreased in C10^G58R^ vs C10^WT^ heart mitochondria, as expected given its autocatalysis when activated (Figure 7A, left). CI and CIV subunits were mildly decreased overall in C10^G58R^ heart mitochondria, consistent with the decreased CI and CIV activities. In contrast to the other CIV subunits, three were increased: COX6A1, COX7A2, and COX7A2L. These CIV isoforms (called the “liver” isoforms) are not typically expressed in adult striated muscle but are the dominant isoforms in most other tissues (47, 48). The “heart” isoforms COX6A2 and COX7A1 were decreased, suggesting that CIV may undergo a subunit switch favoring liver over heart subunits (Figures 7A, B, and E). Enzymes involved in coenzyme Q metabolism were also found to be overall decreased, with the exception of the upstream enzyme PDSS2, similar to what has been previously observed in models of decreased mtDNA expression (19).

**Figure 7.**
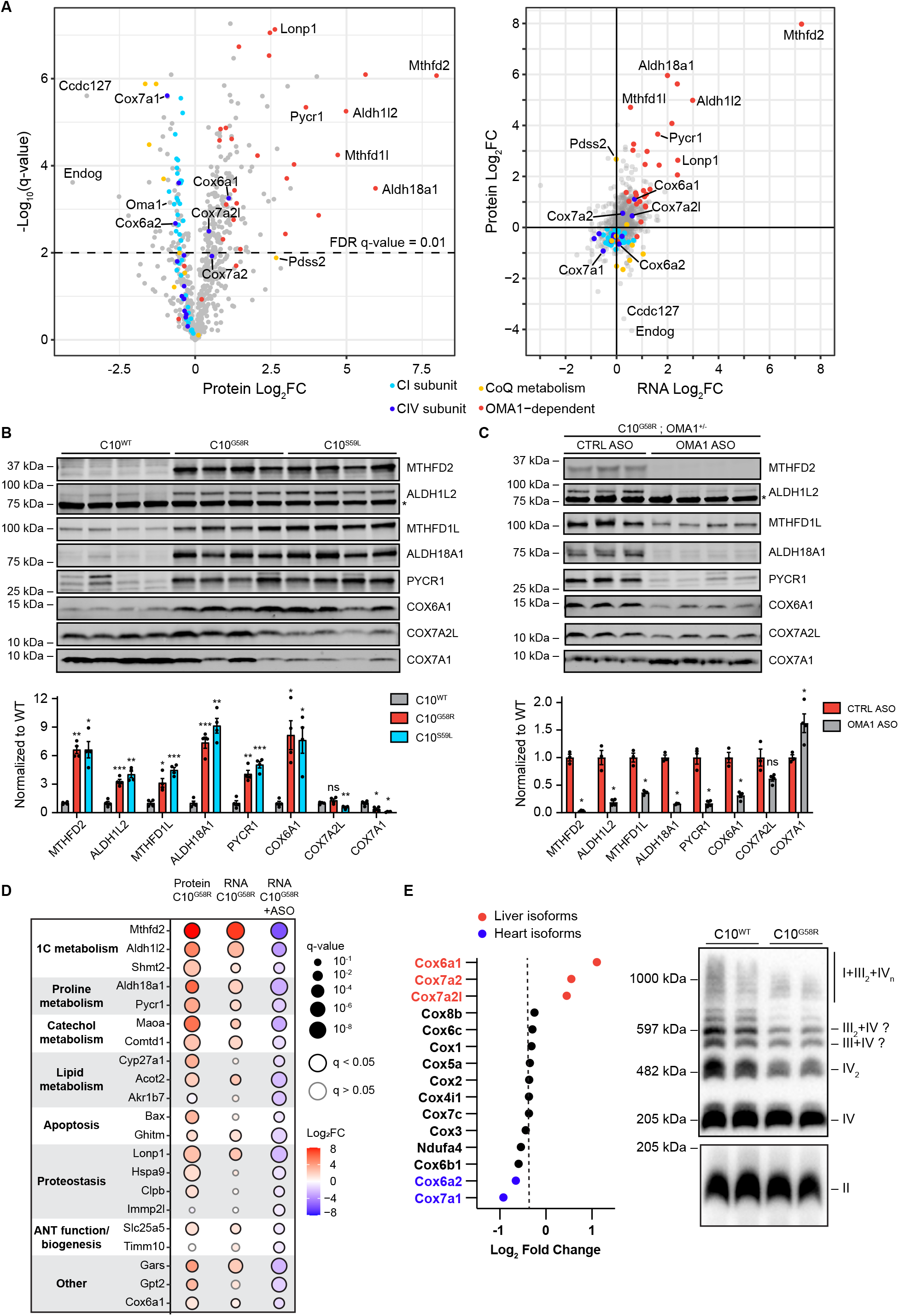
OMA1 activation shapes the mito-proteome through mitonuclear signaling. (A) Left: volcano plot of protein fold change in 36-week-old C10^G58R^ vs C10^WT^ mouse heart mitochondria quantified by label-free mass spectrometry (n = 4 mice per group). Right: protein fold change vs transcript fold change in C10^G58R^ vs C10^WT^ mouse hearts. The proteomics data is 36-week-old C10^G58R^ vs C10^WT^ mice on the WT background, and the transcriptomics is from 14-week-old C10^G58R^ vs C10^WT^ littermates on the OMA1^+/-^ background. (B) Validation of some proteins that were significantly upregulated or downregulated in the C10^G58R^ vs C10^WT^ proteomics dataset. Immunoblot of C10^WT^, C10^G58R^, and C10^S59L^ mouse heart lysates. Loading controls in Figure S7A. *Nonspecific band. (C) Immunoblot of heart lysates from 33-week-old C10^G58R^ mice on the OMA1^+/-^ background treated with a nontargeting (CTRL) or OMA1 ASO. Loading controls in Figure S7B. *Nonspecific band. (D) Comparison of proteomic data from C10^G58R^ vs C10^WT^, transcriptomic data from C10^G58R^ vs C10^WT^ on the OMA1^+/-^ background, and transcriptomic data from OMA1^+/-^ ; C10^G58R^ OMA1 ASO vs CTRL ASO for high-confidence OMA1 targets captured by proteomics. (E) Left: foldchange of complex IV subunits from the C10^G58R^ vs C10^WT^ heart mitochondrial proteomics experiment in (A). Right: blue native PAGE gel of heart mitochondrial lysate of 36-week-old C10^WT^ and C10^G58R^ mice, stained for complex IV (top) and complex II (bottom). Error bars represent SEM. *p<0.05, **p<0.01, ***p<0.001, ****p<0.0001. See also Figure S7 and Supplemental File 4.

Plotting protein vs RNA expression showed almost all OMA1-regulated genes were concordantly increased, suggesting that these protein changes are likely transcriptionally driven (Figure 7A, right). Among the most increased OMA1-dependent genes were enzymes in the mitochondrial one-carbon metabolism and the proline synthesis pathways, which are known targets of the ISR. Notably, the liver COX isoforms were also concordantly increased, whereas the heart isoform COX7A1 was concordantly decreased, suggesting that the COX isoform switch may also be mediated by the ISR.

We next assessed whether these changes to the mitochondrial proteome depended on OMA1 (Figures 7B and S7A). Enzymes in 1C metabolism and proline synthesis pathways were significantly increased in C10^G58R^ and C10^S59L^ hearts, and were suppressed by OMA1 KD in C10^G58R^ hearts (Figures 7C and S7B), confirming that their upregulation at the protein level is signaled by OMA1. Similarly, the liver isoform COX6A1 increased in C10^G58R^ and C10^S59L^ hearts and was suppressed in C10^G58R^ hearts by OMA1 ASO, confirming that it is also upregulated by OMA1. COX7A2L exhibited a similar pattern to COX6A1 but did not reach significance. By contrast, the heart isoform COX7A1 exhibited the converse pattern; it was decreased in both C10^G58R^ and C10^S59L^ hearts and increased in C10^G58R^ hearts treated with OMA1 ASO, suggesting that it may be negatively regulated by OMA1. Considering all identified high-confidence OMA1-regulated mitochondrial genes, the majority were significantly upregulated at both the transcript and protein levels and suppressed by OMA1 ASO at the transcript level (Figure 7D). These results demonstrate that an OMA1-dependent transcriptional response rewires the heart mitochondrial proteome in response to the G58R mutation.

We identified that OMA1 mediates a heart-to-liver isoform switch in CIV of C10^G58R^ mouse hearts. Notably, this switch occurred in each of the intact CIV complexes, separated by blue native PAGE (BN-PAGE) gel electrophoresis (Figures S7C and D, and Supplemental File 4). Overall, the monomeric form of CIV was relatively preserved in C10^G58R^ hearts, but CIV-containing supercomplexes were decreased (Figure 7E). Together, these results demonstrate that the OMA1-dependent ISR reconfigures intact CIV to contain the more ubiquitous CIV liver isoforms.

## Discussion

Through genetic characterization of a C10 G58R family and a novel C10^G58R^ KI mouse model, we have established that the G58R mutation is pathogenic, causing aggregate formation and autosomal dominant isolated mitochondrial myopathy (IMMD). OMA1 is activated in response to C10 G58R misfolding to mediate a protective response that involves mitochondrial fragmentation locally and signaling outside the mitochondria to activate the ISR. The ISR reshapes the mitochondrial proteome, including upregulation of the liver isoforms of CIV subunits, which we identify as a novel component of the ISR in striated muscle. To our knowledge, this is the first demonstration that the OMA1 stress response can be essential for neonatal survival and can mediate mitonuclear signaling through the ISR *in vivo*.

The heterozygous C10 G58R variant was first identified *in cis* with R15S in a large US family with autosomal dominant myopathy (30, 31). Because the two mutations co-segregated, it was not clear which variant was pathogenic. Here, we presented a family from the UK exhibiting myopathy and cardiomyopathy with a *de novo* G58R mutation in isolation, establishing that the G58R variant is pathogenic. The presentations of the US and UK families share several features, including (a) a generalized myopathy presenting in the first decade of life, (b) myopathic features and absence of fibrillations on needle electromyography, (c) moderately decreased complex II-III activities and severely decreased complex IV activity, and (d) intramitochondrial inclusions that we found to be dilations of the cristae with intracristal membranes.

Despite their similarities, the US and UK families differed in disease severity. In the US family some members lived into their 60s, facial weakness was delayed until the third and fourth decade, and cardiomyopathy was not a general feature (31). By contrast, the three affected members of the UK family all died before the age of 35, and all had early facial weakness and cardiomyopathy. Although the reason for this difference is not clear at present, genetic background including a potential protective effect of the *cis* R15S variant may play a role.

The C10 G58R and S59L mutations – while affecting neighboring residues – cause different phenotypes, with a more severe myopathy in G58R patients and upper and lower motor neuron involvement (as well as cerebellar, cortical, and possible nigrostriatal involvement) in S59L patients (32). As in humans, we found myopathy to be more severe in the C10^G58R^ mice than in the C10^S59L^ mice. The differential toxicities of G58R and S59L in muscle and nerve may relate to the distinct toxic conformations they adopt. Notably, C10 G58R and S59L exhibited differential solubility and formed aggregates with distinct morphologies and localizations in the affected heart tissue. Thus, while misfolding into toxic conformations may be a shared pathogenetic mechanism for dominant C10 and C2 mutations, distinct toxic conformations may cause distinct phenotypes. This parallels recent observations of other toxic misfolding proteins, such as α-synuclein and tau, which misfold into different pathogenic strains, each of which causes a distinct but related neurodegenerative disorder (49, 50).

C10 G58R dramatically distorted the IMM, with mitochondrial cristae forming focal dilations with internal vesicles. The IMM is commonly altered in models with mitochondrial dysfunction, and rod-like intracristal inclusions are known to develop in response to creatine depletion (51). However, the morphology of C10 G58R inclusions was distinct and plausibly reflects strain on the IMM from C10 G58R misfolding. Notably, the disruption of the IMM was more pronounced in the C10^G58R^ model than the comparatively mild effects on OXPHOS subunit expression, suggesting that IMM disruption may be a proximal event. We propose that the morphogenesis of these intracristal inclusions may involve dilation of the cristae followed by internalization of membranes (possibly by inward budding), as some dilated cristae were empty but most cristae with internal vesicles were dilated. Interestingly, L-OPA1, which is thought to stabilize the neck of cristae (52), may inhibit the formation of these inclusions, as fewer inclusions formed in the absence of OMA1. It is not clear at present whether C10 G58R disrupts the IMM through a direct interaction with the membrane or through an indirect mechanism.

Strikingly, OMA1 activation was essential for neonatal survival in C10^G58R^ mice. This contrasts with previously-reported models, including YME1L conditional heart KO and PHB2 conditional brain KO, in which OMA1 activation drove the pathology (23, 24). The trigger for OMA1 activation, however, may be different in these models; YME1L, OMA1, and PHB2 associate within inner membrane proteolytic hubs, the disruption of which may lead to aberrant OMA1 activation (53, 54). Activation of OMA1 by C10 G58R, by contrast, appears to be through ΔΨ_m_ instability, likely as a consequence of decreased IMM integrity. This mechanism may be similar to what has been observed in other models, such as Drp1 depletion (42, 43). Both sustained ΔΨ_m_ loss and unstable ΔΨ_m_ are directly sensed by a voltage sensor in OMA1 to increase its activity (11, 42, 43). Thus, in the setting of protein aggregates that disrupt IMM integrity, OMA1 appears to mediate a protective stress response. Notably, C10 G58R, which caused greater IMM distortion than S59L, also activated OMA1 more strongly and was more dependent on OMA1 for survival.

In addition to demonstrating that OMA1 mediates mitochondrial fragmentation in response to C10 G58R stress, we show for the first time *in vivo* that OMA1 signals mitochondrial stress to the nucleus through the ISR. This likely occurs by OMA1 cleavage of DELE1 in the IMS, as part of the OMA1-DELE1-HRI pathway recently described in cell culture studies (12, 13). This mechanism is likely shared by other mitochondrial stresses that disrupt inner membrane integrity or otherwise cause instability of ΔΨ_m_ to activate OMA1.

Our findings beg the question: do all forms of mitochondrial dysfunction activate the ISR through OMA1? Notably, the ISR is potently activated by mutations or toxins that decrease mtDNA expression (sometimes termed mitonuclear imbalance) (19, 22). Are these stress responses also OMA1-dependent? A couple of lines of evidence suggest that they may not be. Although cell culture studies found that diverse mitochondrial toxins activate the ISR through OMA1 and DELE1, inhibition of mtRNA translation with doxycycline did not (13). Additionally, using OMA1 levels as a biomarker of OMA1 activation, we compared the heart mito-proteome of our C10^G58R^ model to models with reduced mtDNA expression (19). Although the ISR was activated to a similar extent in all models, OMA1 levels were reduced as expected in the C10^G58R^ model but unchanged or increased following diminished mtDNA expression (Figure S7E). Notably, models with diminished mtDNA expression had larger reductions in OXPHOS subunits than we observed in our C10^G58R^ model (Figure S7E). We speculate that there are at least two mitochondrial ISR pathways: one that signals loss of IMM integrity through OMA1, and another that signals severe OXPHOS deficiency independently of OMA1. Plausible mechanisms for the OMA1-independent ISR activation include GCN2 activation in response to asparagine deficiency or mTOR activation, as proposed previously (18, 21, 55).

In addition to well-characterized ISR-dependent changes in the heart, we observed upregulation of the ubiquitous liver subunits of CIV (including COX6A1, COX7A2, and COX7A2L), and downregulation of their heart counterparts (COX6A2 and COX7A1). Analysis of proteomics data from a previous study of five models with decreased mtDNA expression showed a similar shift between the liver and heart isoforms, suggesting that this is a general feature of the ISR in striated muscle (Figure S7F) (19). Although the liver isoforms are present in striated muscle at birth in mice, their expression decreases around 4 weeks of age, as they are replaced by their heart counterparts (56). Although COX7A2L has been observed previously to increase in response to ER-stress-mediated ISR (through PERK) (57), our findings place this change in the broader context of a subunit switch in CIV favoring the more generally expressed liver subunits.

In summary, we have shown that the C10 G58R mutation causes a pathologically distinct form of mitochondrial myopathy, IMMD. In response to C10 G58R misfolding, OMA1 mediates a protective stress response which fragments mitochondria locally and signals mitochondrial stress to the nucleus through the ISR. As the OMA1 stress response is protective, interventions aimed at facilitating this response may have therapeutic potential. Additionally, as C10 G58R has a toxic gain of function mechanism and C10 KO is tolerated (20, 27), knockdown of C10 may be a viable therapeutic strategy for IMMD.

## Methods

### Materials Availability

Mouse lines and unique materials generated for this study will be made available upon a reasonable request to the corresponding author.

### Data availability

Microarray data generated in this study has been deposited to GEO (GSE189393 and GSE189396) and the analyzed data is available in Supplemental File 2. Proteomics data is available in Supplemental File 4.

### Genetic characterization of CHCHD10 p.G58R family

The proband of the family was referred to the Mitochondrial Diagnostic Centre at Oxford for clinical assessment, histological, biochemical and molecular genetic analyses.

### Mouse models

Mice were maintained on a 12-hour light/dark cycle, with food and water provided *ad libitum*. The C10^G58R^ mouse was generated using clustered regularly interspaced short palindromic repeats (CRISPR)/caspase 9 (Cas9) endonuclease-mediated genome editing on the C57Bl6J background. A guide RNA (gRNA) near the G54 sequence in mouse C10 (mouse equivalent of G58) was used (TAGCCGTGGGCTCAGCTGTAGGG), along with a single-stranded donor oligonucleotide with a GGC → AGA substitution at G54 for repair: CTGCCGCTCCCGGCCAGCCGGGTCTTATGGCTCAGATGGCATCCACCGCCGCAGGCGTAGCCGTGAGATCAGCTGTAGGCCATGTCATGGGTAGCGCCCTGACCAGTGCCTTCAGTGGGGGAAATTCAGAGCCTGCCCAGCCTGCCGTCCAGCAGGTGAGCGGGAGGACTCAAGAAACGGAGGCAGGATTCACACATGGT. The following primers were used for genotyping: forward primer 5’-GACCCTGGAGTAGAGGGGTT-3’ and reverse primer 5’-GGCCACTCCTCATTGGACTC-3’. The mouse C10 G54R mutation removes a cut site of the restriction endonuclease BanII (NEB, cat# R0119S), introducing a restriction fragment length polymorphism. Therefore, the region surrounding G54 was amplified by PCR and then subjected to a BanII restriction digest for 1 hour followed by 10 minutes of heat inactivation at 80 C, and finally run on a 2% agarose mini gel. The digested WT bands are ∼150bp and the undigested G58R bands are ∼300bp.

The generation of C10^S59L^, C2/C10 DKO, OMA1 KO, and mito-QC mice was described previously (20, 44, 58).

### Motor function tests

C10^WT^ and C10^G58R^ littermates aged 18 weeks and C10^S59L^ mice aged 25 weeks were tested. Mouse forelimb grip strength was measured by pulling the mouse and recording the force generated as the mouse grips to the instrument (BIOSEB instrument with bar, cat# EB1-BIO-GS3). The mouse had to grip with both forelimbs for the trial to be scored; if the mouse used only one forelimb or did not have a proper grip, it was allowed to rest and the trial was repeated. This was repeated 3 times with 15 seconds of rest in between, and the results were averaged.

Balance and motor coordination were tested by placing the mice on a rotating rod (rotarod, Ugo Basile cat# 57624) and measuring the time to fall. The rotarod was allowed to rotate briefly to ensure that all mice faced the proper direction, and when all were facing forwards the speed was ramped from 5 rpm to 40 rpm over 120 seconds; the trial finished when all the mice had fallen off the rotarod. This was repeated 3 times, with 15 minutes of rest between each trial.

Endurance and fatigue were measured with the treadmill fatigue test adapted from a previously-published protocol (59). The mice were run on a treadmill (Columbus Instruments cat# 1050-RM Exer-3/6) with gradually increasing speed, and the distance at which the mouse begins spending most of the time in the rear end of the treadmill (defined as 5 consecutive seconds being within a body’s length of the shocker) was recorded; this was performed after three days of training. The electric stimulus was at 2Hz and 1.125 mA. For training, the mice were run for 5 minutes at a speed of 10 meters/minute. This was done once a day for the three days prior to the test. For the fatigue test, mice were run at the following speeds for the specified amounts of time: 30 seconds at 8 m/min, 30 seconds at 10 m/min, 3 minutes at 12 m/min, 4 minutes at 16 m/min, 4 minutes at 20 m/min, 4 minutes at 24 m/min, and then at 28 m/min until the mouse is fatigued. Mice were determined to be fatigued and removed from the treadmill when they spent 5 consecutive seconds within a body’s length of the shocker, and the distance run was recorded.

As C10^G58R^ mice were generally unable to turn when placed facing upwards in the pole test, the test had to be modified so that mice are placed at the top of the 50 cm pole facing downwards and allowed to descend. To train the mice, this was repeated 3 times with 1 minute of rest in between, and this was done each of the two days preceding the experiment. After the two training days, mice were placed at the top of the pole and the time needed until all four paws touch the base was recorded. This was repeated 5 times with 3 minutes of rest between each trial, and the trial results were averaged.

The behavioral tests were performed in the same week as follows – day 1: grip strength, pole test training 1, treadmill training 1; day 2: rotarod, pole test training 2, treadmill training 2; day 3: pole test, treadmill training 3; day 4: treadmill.

### Echocardiography and electrocardiography

Mouse heart echocardiography and electrocardiography (ECG) were performed at the NHLBI Phenotyping Core. Mice were lightly anesthetized with isoflurane and placed supine over a heated platform with ECG leads and a rectal temperature probe. The Vevo2100 ultrasound system (VisualSonics, Toronto, Canada) with a 30 MHz ultrasound probe (VisualSonics, MS-400 transducer) was used to acquire heart images. Measurements were made from standard 2D and M mode images from the parasternal long axis and mid-papillary short axis views of the left ventricle.

### Mouse complexes I and IV activity assays

Assays were performed according to the kit instructions (Abcam CI rodent: AB109721, CIV rodent: AB109911). All preparation steps, including homogenization, were performed on ice, and 30-60 mg samples of liquid-nitrogen-flash-frozen mouse heart or muscle were used.

### Mouse mtDNA long-range PCR

Whole DNA was extracted from 15-25 mg of mouse tissue using the DNeasy Blood and Tissue Kit (QIAGEN, cat# 69504) according to the kit’s instructions. DNA was then diluted in water to 40 ng/μL. The following primers were used – forward: 5’-CTGGAATTCAGCCTACTAGCAATTATCC-3’ and reverse: 5’-TTTAGGTTTATGGCTAAGCATAGTGGGG-3’ (60). The final PCR reaction contained 10 μL of Platinum SuperFi II Green Master Mix (Invitrogen, cat# 12369010), 5μL template DNA (40 ng/μL, 200 ng total), and 5 μL of primer master mix (1 μM forward and 1 μM reverse primers in water, for a final reaction concentration of 250 nM each). The PCR was run with the following cycle: 98 C for 2 minutes, 20 x (98 C for 10 seconds, 60 C for 15 seconds, 72 C for 6 minutes and 24 seconds), 10 x (98 C for 10 seconds, 60 C for 15 seconds, 72 C for 6 minutes and 24 seconds + 20 seconds per cycle), 72 C for 5 minutes, 4 C indefinitely.

Following the PCR, the 20μL samples were diluted in 40 μL of water and run on a 0.8% agarose gel for 1 hour at 70 Volts in TAE buffer and post-stained with SYBR Gold (Invitrogen, cat# S11494) for 30 minutes in TAE buffer. Gels were imaged on a Bio-Rad Chemidoc system.

### Mouse mtDNA copy number

Whole DNA was extracted from hearts as described above. DNA was digested with HaeIII (NEB, cat# R0108T) as follows: 5 μL 10X CutSmart buffer, 1 μL HaeIII, 500 ng DNA, and water for a final reaction volume of 50 μL at 37 C and 300 RPM for 15 minutes, followed by a 5-minute heat inactivation at 80 C. DNA was diluted in water from 50 ng/μL to 1 ng/μL. Droplet digital PCR (ddPCR) primer/probe assays for copy number variation (CNV) were obtained from Bio-Rad (mouse ND1/HEX, mouse ActB/FAM). The ddPCR mix was as follows: 3.5 μL digested DNA (1 ng/μL), 1.1 μL ND1 assay, 1.1 μL ActB assay, 5.3 μL water, 11 μL 2X ddPCR Supermix for Probes. Droplets were generated with the Automated Droplet Generator (Bio-Rad, cat# 1864101), and the reaction was run with CNV settings using the QX200 Droplet Reader (Bio-Rad, cat# 1864003). The resulting data was analyzed with QuantaSoft Analysis Pro (Bio-Rad).

### Human mtDNA long-range PCR

Genomic DNA was extracted from fresh frozen tissue by first homogenizing tissue and then using conventional proteinase K digestion and phenol-chloroform extraction methods. Long range PCR of 13.8 kb of the human mitochondrial genome (NC_012920.1 m.2695 to m.16459) was undertaken using primers described by Li et al., 1995 (named L1 and H3) and TaKaRa LA Taq HS (Takara Bio Inc., Japan), followed by agarose gel electrophoresis (61).

### Human histochemistry and complex activities

Histochemistry and ETC complex activities were performed according to standard methods within the context of an accredited NHS diagnostic laboratory (62–64).

### Mouse COX activity stain and fiber width measurements

Mice were anesthetized with isoflurane and transcardially perfused with PBS. Tibialis and soleus muscles were collected and immediately flash-frozen in liquid-nitrogen-cooled isopentane and stored at −80 C until sectioning. The muscles were cryosectioned at −20 C to obtain 10 μm transverse sections. For staining, muscle sections were allowed to dry at room temperature for 1 hour and then were incubated in the incubation solution for 1 hour at 37 C. Incubation solution: 100 μM cytochrome c (Sigma, cat# C2506), 4 mM DAB (Sigma, cat# D5637), 20 μg/mL catalase (Sigma, cat# C40) in 0.2 M phosphate buffer (pH 7.0). Sections were washed in 0.2 M phosphate buffer for 5 minutes twice, mounted with mounting medium (KPL, cat# 71-00-16), coverslipped, and imaged with a Zeiss Wide-Field microscope.

Image analysis was performed in Fiji (NIH). Muscle fiber maximal widths were measured for 20 random fibers per mouse.

### Lipid staining

Mice were anesthetized with isoflurane and transcardially perfused with 20 mL PBS followed by 15 mL of 4% paraformaldehyde (PFA). Tibialis muscles were dissected and postfixed in 4% PFA for 30 minutes at room temperature, washed for 5 minutes in PBS 3 times, and stored in PBS at 4 C until use. Longitudinal 35 μm free-floating sections were obtained using the Compresstome VF-300-0Z (Precisionary). Muscle sections were incubated in 1:1000 LipidTOX deep red (Thermo Fisher Scientific, cat# H34477) for 1 hour at room temperature on a rocker and protected from light. The sections were then washed with PBS for 10 minutes 3 times, mounted on slides with mounting medium (KPL, cat# 71-00-16), coverslipped with 1.5H thickness coverslips (Thorlabs, cat# CG15KH), and sealed with nail polish. Sections were imaged on an Olympus FLUOVIEW FV3000 confocal laser scanning microscope.

### Indirect calorimetry

The Oxymax-CLAMS setup (Columbus Instruments) was used to assess mouse metabolism. Mice were singly housed in the CLAMS chambers, received food and water *ad libitum* throughout the experiment, and were checked on at least twice a day throughout the 5-day duration of the experiment. Oxygen consumption, carbon dioxide production, food intake, and beam breaks (locomotion) were monitored during the experiment. The resulting data was analyzed using CalR (65).

### Body composition

Non-invasive measurements of lean tissue and fat were obtained using the EchoMRI NMR machine. Un-anesthetized mice were placed in a clear plastic tube and gently restrained at the end of the tube by using a plunger with air holes. The plunger was fitted to the mouse and tightened gently to minimize movement. The tube was then inserted into the mini-spec port to a premeasured depth, and measurements were collected. The mouse was removed from the tube and returned to the home cage, and the tube was washed and sanitized after each use.

### EVcouplings conservation analysis and 3D prediction

The human CHCHD10 protein sequence (Q8WYQ3) was analyzed using the EVcouplings server (https://v2.evcouplings.org/). The b0.2 cutoff yielded the most matched sequences and was used in the subsequent analysis. Outputs included assessment of amino acid frequency for each residue among the identified C10 homologs, as well as structure predictions generated using the EVfold algorithm (40, 41). The highest scoring structure was visualized using UCSF Chimera (https://www.cgl.ucsf.edu/chimera/).

### Hydrophobicity analysis

Hydrophobicity of the middle a-helix of WT, G58R, and S59L C10 was calculated using ProtScale and the Kyte-Doolittle algorithm (https://web.expasy.org/protscale/).

### Cell transfection

HeLa cells were transfected with WT or mutant C10 constructs using the FuGENE 6 transfection reagent (Promega, cat# PRE2691). The cells were incubated at 37 C at least overnight before downstream analysis.

### Immunoblotting

Immunoblotting and densitometry measurements were performed as described previously (20).

### Mitochondrial isolation

Mitochondria were isolated from whole mouse hearts as described previously (66).

### Soluble/insoluble assay

This assay was performed on cultured cells or on mouse tissue. HEK293 or HeLa cells were harvested by scraping into PBS 24 hours after transfection, whereas mouse tissue was fractionated and the assay was performed on total homogenate, the cytosolic fraction, and the mitochondrial fraction. An aliquot was used to determine protein concentration using the BCA kit, and normalization was performed according to protein content. The cells were pelleted at 400 g and resuspended in Triton X-100 lysis buffer (20 mM Tris-HCl pH 7.5, 150 mM NaCl, 10 mM EDTA pH 8.0, 1% Na-DOC, 1% Triton X-100). Afterwards, the sample was centrifuged at 21130 g to obtain soluble (supernatant) and insoluble (pellet) fractions. Laemmli buffer was added to both fractions to obtain an equal volume of 1X buffer with 2.5% (v/v) 2-mercaptoethanol. Lysates were boiled at 98 C, separated by SDS-PAGE, and analyzed by immunoblotting.

### Mouse histochemistry

Mice were anesthetized with isoflurane and transcardially perfused with 25 mL PBS. The heart midventricular region and tibialis muscle were dissected and postfixed in 4% PFA for 1 hour (heart) or 30 minutes (tibialis) at room temperature, with the heart additionally fixed overnight at 4 C. After fixation, the tissue was washed for 5 minutes in PBS 3 times. Some heart specimens were sent to Histoserv (Germantown, MD) for paraffin embedding and microtome sectioning to obtain 4 μm sections. H&E and Masson’s trichrome stains were additionally performed by Histoserv.

35 μm free-floating sections of heart and muscle were obtained using the Compresstome VF-300-0Z (Precisionary). The sections were permeabilized and blocked in 0.4% Triton-X (Sigma, X100) and 4% BSA fraction V (MP Biomedicals, cat# 02160069) in PBS for 2 hours at room temperature. Afterwards, the sections underwent primary antibody incubation in 0.3% Triton-X and 1% BSA in PBS overnight at 4 C (note: samples from the PDH/C10 experiment from Figure 2G were incubated in primary for 3 days). The samples were washed for 10 minutes in PBS 3 times, and then incubated in secondary antibodies in PBS (1:500). Afterwards, the samples were washed for 10 minutes in PBS 3 times, mounted on a slide with mounting medium (KPL, cat# 71-00-16), coverslipped (Thorlabs, cat# CG15KH), and sealed with nail polish.

The p62/C10 staining from Figure S4E was performed on unstained 4 μm paraffin-embedded heart sections obtained from Histoserv. The staining procedure was identical to the above, but the slides had to be deparaffinized first with the following serial incubations: 2×5 minutes xylenes (Sigma, cat# 534056) twice, 3 minutes 1:1 xylenes:ethanol, 2×3 minutes 100% ethanol, 3 minutes 95% ethanol, 3 minutes 70% ethanol, 3 minutes 50% ethanol, 30 seconds rinse in tap water.

### Super-resolution imaging

The super-resolution images shown in Fig. 2D, 2G, 2H, and 5A-C were acquired using a LSM 880 with Airyscan (Zeiss) microscope as z-stacks with a minimum of 5 images. The images were 3D-deconvolved using the default settings in the Zeiss Zen software. The images shown are maximum intensity projections of the z-stack produced using ImageJ/Fiji (NIH).

### Immunogold-electron microscopy

The method was adapted from a previously-published protocol (67). Briefly, DOX-inducible HEK293 cells were treated with 1 μg/mL doxycycline overnight and then fixed with 4% paraformaldehyde + 0.05% glutaraldehyde for 45 minutes in PBS. The sample was then blocked and made permeable with 0.1% saponin with 5% bovine serum albumin (BSA) for 40-60 minutes. The sample was incubated with CHCHD10 (Sigma, cat# HPA003440) primary antibody followed by secondary antibodies (Nanogold, Nanoprobes, Yaphank, NY) for 1-2 hours, postfixed with 2% glutaraldehyde in PBS for at least overnight, silver enhanced (HQ kit, Nanoprobes), *en bloc* mordanted with 0.25-0.5% uranyl acetate in acetate buffer (pH 5.0) for 1 hour, treated with 0.2% osmium tetroxide in 0.1 M phosphate buffer (pH 7.4) for 30 minutes, dehydrated with a series of graded ethanol, and embedded in epoxy resin. Imaging was performed on a JEOL-1200 EXII TEM.

### Transmission electron microscopy

Human pectoralis muscle was snap frozen in liquid-nitrogen-cooled isopentane and subsequently processed for electron microscopy according to standard protocols.

Mouse heart and skeletal muscle TEM was performed as previously described (20).For the serial TEM sections, the samples were prepared with a modified protocol for heavy membrane staining. Heart samples were obtained and fixed with glutaraldehyde as above. The samples were placed in filtered 1% tannic acid solution in EM buffer on ice for 1 hour. The samples were then washed and placed in a solution of 1% osmium tetroxide and 1% ferrocyanide in EM buffer. After that, the samples were washed and processed *en bloc* as above with uranyl acetate, dehydration, embedding, serial sectioning, and imaging.

### Assessment of cell lines

HeLa^OMA1 KO^ cells were a gift from Richard Youle (National Institutes of Health) and their generation was described previously (68). HEK293^C2/C10 DKO^ and HEK293 C10 G58R Tet-inducible cells were described previously (20). To assess mitochondrial potential stability, Tet-inducible HEK293 C10 G58R cells also expressing mito-mEmerald were treated with DMSO or 1 μg/mL doxycycline overnight. The cells were then stained with 5 nM TMRE and imaged live using an Olympus FLUOVIEW FV3000 confocal microscope in resonance mode to visualize membrane potential fluctuations. Images were obtained every ∼500 msec for 90 seconds. Cells were scored as having ΔΨ_m_ instability if the TMRE intensity of a mitochondrion within the field was observed to decrease and then increase again during the 90 second visualization period, in the setting of a constant mito-mEmerald signal. To assess ΔΨ_m_ by flow cytometry, Tet-inducible HEK293 C10 G58R cells were treated overnight with DMSO or 1 μg/mL doxycycline. The cells were treated with 10 μM CCCP or 10 μM oligomycin together with 5 nM TMRE for 10 minutes, and their membrane potential was measured using a CellStream (Luminex) flow cytometer. All cell lines were incubated at 37 C with 5% CO_2_ and ambient O_2_.

### FIB-SEM, segmentation, reconstruction, and analysis

Heart samples were processed with the standard TEM protocol described above. The blocks were processed with the ZEISS Crossbeam 540 (Gemini II column) to collect FIB-SEM micrographs at a 10 nm pixel size (X x Y x Z) with the ZEISS Atlas 5 software (Carl Zeiss Microscopy GmbH, Jena, Germany). The 10 nm thickness FIB milling was conducted at 30 keV while maintaining a beam current at 2 nA. The collected micrographs were aligned using a proprietary algorithm and then exported as 8-bit TIFF files, which were sent to Ariadne (https://ariadne.ai/, Switzerland) for AI-based segmentation of mitochondria. The 3D-reconstructed mitochondria for this paper were generated using Dragonfly software, version 2020.2 for Windows 10 (Object Research Systems Inc, Montreal, Canada, 2020; software available at http://www.theobjects.com/dragonfly). Analysis was performed using the multi-ROI object analysis tool in Dragonfly.

### Microarray and analysis

RNA was extracted from mouse hearts using either the Direct-zol RNA Miniprep kit (Zymo, cat# R2051) or the RNeasy Fibrous Tissue Mini Kit (QIAGEN, cat# 74704, used for the ASO experiment). RNA expression was measured using the Clariom_S_Mouse microarray (Affymetrix).The Transcriptome Analysis Console software (Affymetrix, version 4.0.1) was used to analyze the data with the default settings. Specifically, data were summarized using the gene-level signal space transformation - robust multiple-array average normalization (SST-RMA) method. DEGs were required to have a genome-wide q-value of < 0.01 or 0.05 (as specified) measured using the eBayes analysis of variance (ANOVA) method. Heatmaps were generated using the ComplexHeatmap package in R. The genes displayed in the heatmaps consisted of ATF4, other transcription factors involved in the ISR, ATF4 target genes, and myc (22, 69–72). GSEA was performed using the GSEA software and reactome pathways (73–75). A separate GSEA was performed using reactome pathways plus a custom “G58R Upregulated” category of genes that were upregulated with an FDR q-value < 0.01 in the C10^G58R^ ; OMA1^+/-^ vs C10^WT^ ; OMA1^+/-^ comparison.

### ASO experiment

ASOs were synthesized at Ionis Pharmaceuticals as previously described (76). C10^G58R^ ; OMA1^+/-^ mice were given weekly subcutaneous injections of either a nontargeting (control) ASO or one of two OMA1-targeting ASOs starting at 21 weeks of age. The ASOs were at a concentration of 5 mg/mL, and injections were dosed at 50 mpk. A total of 12 injections were administered per mouse over 12 weeks. Echocardiography was performed during the 10^th^ week, and motor function tests were performed during the 12^th^ (final) week. After the tests were completed, the mice were anesthetized with isoflurane, transcardially perfused with PBS, and their tissue was collected and flash frozen in liquid nitrogen. Immunoblotting and transcriptomics were performed on heart tissue as described above.

### Proteomics and analysis

Mitochondria were isolated from newly harvested mouse hearts as described above. Protein concentration of the mitochondrial fraction was determined using a BCA kit. 86 μg of the mitochondrial fraction was solubilized in 1% digitonin and complexes were separated on a blue native (BN)-PAGE gel with bovine heart mitochondria used as a molecular weight standard. One gel band was cut for each lane. In-gel proteins were reduced with 10 mM Tris(2-carboxyethyl)phosphine hydrochloride (TECP), alkylated with 10 mM N-Ethylmaleimide (NEM), and digested with trypsin. The extracted peptides were desalted and used for label-free quantitation. Liquid chromatography-tandem mass spectrometry (LC-MS/MS) data acquisition was performed on an Orbitrap Lumos mass spectrometer (Thermo Fisher Scientific) coupled with a 3000 Ultimate high-pressure liquid chromatography instrument (Thermo Fisher Scientific). Peptides were separated on an ES802 column (Thermo Fisher Scientific) with the mobile phase B (0.1% formic acid in acetonitrile) increasing from 3 to 24% over 95 minutes. The LC-MS/MS data were acquired in data-dependent mode. For the survey scan, the mass range was 400-1500 m/z; the resolution was 120 k; the automatic gain control (AGC) value was 8e5. The MS1 cycle time was set to 3 sec. As many MS2 scans as possible were acquired within the cycle time. MS2 scans were acquired in ion-trap with an isolation window of 1.6 Da. Database search and mutant/WT ratio calculation were performed using Proteome Discoverer 2.4 (Thermo Fisher Scientific) against Sprot Mouse database. Proteins were annotated as mitochondrial if they appeared in mouse MitoCarta3.0, and were additionally annotated with MitoPathways from MitoCarta3.0 (77). Identified proteins tagged as non-mitochondrial or without quantifications in at least 3 (Figure 7A) or 2 (Figure S7D) samples in at least 1 group were filtered out using Perseus (MaxQuant).

For the complex IV monomer vs supercomplexes experiment, the BN-PAGE gel was cut into 4 slices as in Figure S7C and proteomics analysis was performed on each slice as above.

### BN-PAGE

Mitochondrial isolation from fresh heart tissue was performed as described above, and BN-PAGE was performed as previously described (28).

### Statistics

FDR q-values were calculated for the transcriptomics experiments with the Transcriptome Analysis Console software (Affymetrix, version 4.0.1), and for the proteomics experiments with Perseus (MaxQuant). All other statistical analyses were performed in GraphPad Prism 9 using Welch’s t test corrected for multiple comparisons, one-way ANOVA with Dunnett’s multiple comparisons, or the log-rank (Mantel-Cox) test for survival data.

### Study approval

The patient research protocol (REC 20/04, IRAS 162181) was approved and performed under the ethical guidelines issued by the South Central-Berkshire Research Ethics committee for clinical studies, with written informed consent obtained for all subjects including for the publishing of patient photographs. All animal studies were approved by the Animal Care Use Committee at the National Institute of Neurological Disorders and Stroke (NINDS) intramural program.

## Author contributions

Conceptualization, M.K.S. and D.P.N.; Methodology, M.K.S. and D.P.N.; Investigation, M.K.S., X.H., B.P.W., I.S., Y.L., C.K.E.B., D.A.S., C.F., I.A.B., J.P., and D.P.N.; Formal Analysis, M.K.S., N.R., D.P.N; Resources, A.F.P., P.M.Q., C.L.O., J.P., and D.P.N.; Writing – Original Draft, M.K.S. and D.P.N.; Writing – Review and Editing, J.P., P.M.Q., M.K.S., and D.P.N.; Visualization, M.K.S. and D.P.N.; Supervision, J.P. and D.P.N.; Funding, J.P. and D.P.N.

## Acknowledgements

We thank Sandra Lara, Dr. Jung-Hwa Tao-Cheng, and the NINDS EM Facility for technical assistance with TEM. We thank Dr. Vincent Schram, Dr. Carolyn Smith, and the NINDS and NICHD Light Microscopy Facilities for technical assistance with brightfield and confocal microscopy. We thank Dr. Abdel Elkahloun and the NHGRI/DIR Microarray Core for technical assistance with RNA expression studies. We thank Steve Shema and the NCI Genomics Core for help with ddPCR. We thank the NINDS Protein/Peptide Sequencing Facility for the label-free quantitative proteomics data acquisition. We thank Dr. Chengyu Liu and the NHLBI Transgenic Core for assistance in generating transgenic mice. We thank Kate Sergeant, Charu Deshpande, and Michael Simpson for their involvement in whole exome sequencing and family testing of the patient, which was supported by the Lily Foundation. Monika Hofer, M Squier and M Esiri carried out histochemical and EM investigations of the family, and Dr. Iain Hargreaves performed respiratory chain studies. We thank Eric Lindberg for his electron microscopy technical expertise. We thank Ionis Pharmaceuticals for providing control and OMA1 targeted ASOs that were used in *in vivo* experiments. We thank Dr. Lucy Forrest for suggestions on structural modeling of CHCHD10. We thank Dr. Richard Youle for critical reading of the manuscript and insightful comments. This work was supported by the Intramural Research Program of the NINDS, National Institutes of Health.

