## Supplemental Figures and Information for "OMA1 mediates local and global stress responses against protein misfolding in CHCHD10 mitochondrial myopathy"

**Resources table**

| REAGENT or RESOURCE | SOURCE | IDENTIFIER |
|---|---|---|
| Antibodies | | |
| CHCHD2 (WB) | Proteintech | Cat#19424-1-AP; RRID: AB_10638907 |
| CHCHD10 (WB, IF 1:250) | Sigma | Cat#HPA003440; RRID: AB_1078348 |
| FLAG M2 (IF 1:1000) | Sigma | Cat#F1804-1MG; RRID: AB_262044 |
| TOM20 F-10 (IF 1:1000) | Santa Cruz | Cat#SC-17764; RRID: AB_628381 |
| Tubulin (WB) | Sigma | Cat#T8328; RRID: AB_1844090 |
| NDUFB8 (WB) | Abcam | Cat#Ab110242; RRID: AB_2756818 |
| PDH (IF 1:100) | Abcam | Cat#AB110333; RRID: AB_10862029 |
| OPA1 (WB) | BD Bioscience | Cat#612606; RRID: AB_399888 |
| OMA1 (WB) | Santa Cruz | Cat#SC-515788 |
| HSP90 (WB) | Proteintech | Cat#13171-1-AP; RRID: AB_2120924 |
| eIF2α (WB) | Santa Cruz | Cat#SC-133132; RRID: AB_1562699 |
| p-eIF2α (WB) | Cell Signaling | Cat#3597; RRID: AB_390740 |
| MTHFD2 (WB) | Proteintech | Cat#12270-1-AP; RRID: AB_2147525 |
| ALDH1L2 (WB) | Proteintech | Cat#21391-1-AP; RRID: AB_2878854 |
| MTHFD1L (WB) | Proteintech | Cat#16113-1-AP; RRID: AB_2250974 |

| | | |
|---|---|---|
| ALDH18A1 (WB) | Proteintech | Cat#17719-1-AP; RRID: AB_2223896 |
| PYCR1 (WB) | Proteintech | Cat#13108-1-AP; RRID: AB_2174878 |
| COX6A1 (WB) | Abcam | Cat#Ab110265; RRID: AB_10866496 |
| COX7A1 | Proteintech | Cat#11413-1-AP; RRID: AB_2085705 |
| COX7A2L (WB) | Proteintech | Cat#11416-1-AP; RRID: AB_2245402 |
| SDHA (BN PAGE) | Abcam | Cat#Ab14715; RRID: AB_301433 |
| MTCO1 (BN PAGE) | Invitrogen | Cat#459600; RRID: AB_2532240 |
| p62 (IF, 1:200) | Abcam | Cat#AB56416; RRID: AB_945626 |
| Chemicals, peptides, and recombinant proteins | | |
| BanII Endonuclease | NEB | Cat#R0119S |
| Platinum SuperFi II Green Master Mix | Invitrogen | Cat#12369010 |
| SYBR Gold | Invitrogen | Cat#S11494 |
| TaKaRa LA Taq Hot Start | Takara Bio | Cat#RR042A |
| HaeIII Endonuclease | NEB | Cat#R0108T |
| Cytochrome c from equine heart | Sigma | Cat#C2506 |
| DAB | Sigma | Cat#D5637 |
| Catalase | Sigma | Cat#C40 |
| Mounting medium | KPL | Cat#71-00-16 |
| LipidTOX deep red | Thermo Fisher Scientific | Cat#H34477 |
| FuGENE 6 | Promega | Cat#PRE2691 |
| BSA fraction V | MP Biomedicals | Cat#02160069 |
| Xylenes | Sigma | Cat#534056 |
| 8% glutaraldehyde | Electron Microscopy Services | Cat#16020 |
| Critical commercial assays | | |
| BCA kit | Thermo Fisher Scientific | Cat#23225 |

| | | |
|---|---|---|
| DNeasy Blood and Tissue Kit | QIAGEN | Cat#69504 |
| Direct-zol RNA Miniprep Kit | Zymo | Cat#R2051 |
| RNeasy Fibrous Tissue Mini Kit | QIAGEN | Cat#74704 |
| Deposited data | | |
| Congenital OMA1 KO vs het effect on G58R and S59L microarray | This paper | GEO: GSE189393 |
| Acute OMA1 KD effect on gene expression in the G58R background microarray | This paper | GEO: GSE189396 |
| Experimental models: Cell lines | | |
| HeLa$^{OMA1\ KO}$ | Sekine et al., 2019 | N/A |
| HEK293$^{C2/C10\ DKO}$ | Liu et al., 2020 | N/A |
| HEK293 C10 G58R Tet-inducible | Liu et al., 2020 | N/A |
| Experimental models: Organisms/strains | | |
| CHCHD10$^{G58R}$ mouse | This paper | N/A |
| CHCHD10$^{S59L}$ mouse | Liu et al., 2020 | N/A |
| CHCHD2/CHCHD10 DKO mouse | Liu et al., 2020 | N/A |
| OMA1 KO mouse | Quiros et al., 2012 | N/A |
| Mito-QC mouse | McWilliams et al., 2016 | N/A |
| Software and algorithms | | |
| CalR | Mina et al., 2018 | RRID: SCR_015849 |
| Fiji | NIH | RRID: SCR_002285 |
| Perseus | MaxQuant | RRID: SCR_015753 |
| Transcriptome Analysis Console Software | Affymetrix | RRID:SCR_018718 |
| Dragonfly | ORS | |
| Image Studio | LI-COR | RRID:SCR_015795 |
| Prism 9 | GraphPad | RRID:SCR_002798 |
| Other | | |
| Grip strength instrument | BIOSEB | Cat#EB1-BIO-GS3 |
| Rotarod | Ugo Basile | Cat#57624 |
| Treadmill | Columbus Instruments | Cat#1050-RM Exer-3/6 |
| Automated droplet generator | Bio-Rad | Cat#1864101 |
| QX200 droplet reader | Bio-Rad | Cat#1864003 |
| 1.5H coverslips | Thorlabs | Cat#CG15KH |

**Supplemental Files and Videos**

Supplemental File 1: Echocardiography of mouse hearts

Supplemental File 2: Gene expression foldchanges of mouse hearts

Supplemental File 3: Gene set enrichment analyses of gene expression data

Supplemental File 4: Proteomics of mouse hearts

Supplemental Video 1: Mitochondrial potential of DMSO- and DOX (C10 G58R overexpressing)-treated HEK293 cells

Supplemental Video 2: FIB-SEM segmentation of heart mitochondria

Supplemental Video 3: Segmented non-megamitochondria (shades of cyan) and megamitochondria (differently colored) from the $C10^{G58R}$ ; $OMA1^{-/-}$ dataset

**Supplemental methods**

*Mouse complexes I and IV activity assays*

30-60 mg samples of liquid-nitrogen-flash-frozen mouse heart or muscle were placed in 1.5 mL centrifuge tubes with 400 µL of Solution 1 from the CIV activity kit. The tissue was minced with scissors into small pieces and then homogenized with a Fisher Scientific PowerGen 125 Homogenizer. Heart was homogenized at power level 4.5 for 1 minute, then 10 seconds at power level 5.5. Muscle was homogenized at power level 4.5 for 1 minute, then 30 seconds at power level 5.5. Protein concentration was determined with a BCA kit (Thermo Fisher Scientific, cat# 23225), and the sample was diluted to 5.5 mg/mL in Solution 1. Detergent was added to the samples for a final volume ratio of 1:10 (e.g. 10 µL Detergent for 90 µL of sample), resulting in a protein concentration of 5 mg/mL. The samples were incubated on ice for 30 minutes then centrifuged at 15,000 g for 20 minutes at 4 C, and the supernatant was collected.

The plates were equilibrated to room temperature before being loaded with samples. For CI activity, heart samples were diluted in incubation solution to 33.75 µg /450 µL, and muscle samples were diluted in incubation solution to 90 µg /450 µL. For CIV activity, heart and muscle samples were diluted to 45 µg/450 µL in Solution 1. 200 µL of diluted sample was added to the respective plate, and the assay was performed in technical duplicates. The plates were incubated at room temperature for 3 hours. Afterwards, wells were emptied and washed thrice with wash buffer (CI) or Solution I (CIV).

For CI, 200 µL of 1X NADH and 1X dye in 1X wash buffer was added to each well, and the plate was placed in a plate reader with the following settings – wavelength: 450 nm, time: 30 minutes, interval: 30 seconds, temperature: room temperature, shake between readings. The slope of the resulting curve represented CI activity.

For CIV, 200 µL of 1:20 reagent c in Solution 1 was added to each well, and the plate was placed in a plate reader with the following settings – wavelength: 550 nm, time: 120 minutes, interval: 1 minute, temperature: 30 C, shake between readings. The absolute value of the slope of the resulting curve represented CIV activity.

*Human mtDNA long-range PCR*

Genomic DNA was extracted from fresh frozen tissue by first homogenizing tissue and then using conventional proteinase K digestion and phenol-chloroform extraction methods. Long-range PCR of 13.8 kb of the human mitochondrial genome (NC_012920.1 m.2695 to m.16459) was undertaken using primers 5'-GAGGCGGGCATAACACAGCAAGACGA-3' and 5'-GGCCCGGAGCGAGGAGAGTAGCAC-3' and TaKaRa LA Taq HS (Takara Bio Inc., cat# RR042A). The reaction conditions were as recommended by Takara Bio for a 25 μL reaction, but with 4 μL TaKaRa dNTP mixture, primers at a final concentration of 1 μM, and approximately 100 ng of template genomic DNA. The PCR was run with the following cycle: 94 C for 1.5 minutes, 25 x (98 C for 10 seconds, 68 C for 10 minutes + 30 seconds per cycle), 72 C for 10 minutes, 15 C indefinitely. Following the PCR, 10 μL of product was run on a 0.7% agarose gel for 4 hours at 80 Volts. Gels were imaged using a UV transilluminator.

*Immunoblotting*

HEK293 and HeLa cells were lysed in a buffer containing 1% sodium dodecyl sulfate (SDS) and 63 mM Tris (pH 6.95). DNA shearing was accomplished using a bath sonicator. Insoluble material was separated by centrifugation at 21130 g at 4 C for 3 minutes and the soluble fraction was retained. Protein concentration was measured using the BCA assay. Bromophenol blue (final concentration 0.0008%), glycerol (11%) and β-mercaptoethanol (0.284 mM) were added to the supernatant. The sample was heated at 80 C for 10 minutes. Insoluble material was separated by centrifugation at 21130 g at room temperature for 3 minutes and the soluble fraction was retained. Subsequently, the soluble fraction was analyzed by sodium dodecyl sulfate-polyacrylamide gel electrophoresis (SDS-PAGE) and immunoblotting.

Mouse tissue was lysed in a buffer containing 20 mM Tris pH 7.8, 137 mM NaCl, 2.7 mM KCl, 1 mM $MgCl_2$, 1% Triton X-100, 10% glycerol, 1 mM ethylenediaminetetraacetic acid, and 1 mM dithiothreitol, with 1% proteinase inhibitor. Lysates were sonicated 4 times by a Vibra-Cell Ultrasonic Disruptor for 15 seconds each time, at an output level of 20. Protein concentration was determined by the BCA assay. Lysates were separated on SDS-PAGE gels and analyzed by immunoblotting.

Densitometry measurements were made by using Fiji (NIH) and Image Studio (LI-COR). OMA1-cleaved S-OPA1 bands were calculated by measuring the maximum intensity of each of the five bands in a linescan of the optical density of the five OPA1 bands on the blot using Fiji (NIH). After subtracting background intensity, the peaks of the c and e bands were summed and divided by the sum of the five bands (a–e) to obtain the percentage of OMA1-generated S-OPA1 from total OPA1.

*Mitochondrial isolation*

The method was adapted from a previously-published protocol (66). Mice were anesthetized with isoflurane, transcardially perfused with 25 mL PBS, and the heart was dissected and placed in a dish with cold PBS. The heart was then transferred into a tube with 10 mM EDTA in PBS and was minced into small pieces with scissors. The sample was centrifuged for 30 seconds at 5000 g and was washed twice by PBS/EDTA solution. After the washes, PBS/10 mM EDTA/0.05% trypsin solution was added and the tube was placed in a rocker for 30 minutes at 4 C. The tube was centrifuged at 200 g for 5 minutes and the supernatant was discarded. The tissue was weighed and 10 times the volume of the IBm1 buffer (6.7 mL of 1 M sucrose, 5 mL of 1 M KCl, 5 mL of 1 M Tris/HCl, 1 mL of 1 M EDTA, 2 mL of 10% BSA with water to make 100 mL of buffer, pH 7.4) was added before tissue was homogenized. The homogenate was centrifuged at 700 g for 10 minutes and the supernatant was collected, and it was then centrifuged at 8000 g for 10 minutes. The cytosolic fraction (supernatant) was discarded and the pellet was resuspended in the IBm2 buffer (25 mL of 1 M sucrose, 1 mL of 1 M Tris/HCl, 3 mL of 0.1 M EGTA/Tris with water to make 100 mL of buffer, pH 7.4) and centrifuged again at 8000 g for 10 minutes. The pellet was washed once more in IBm2 buffer before it was resuspended in RIPA buffer with protease inhibitor and phosphatase inhibitor. The sample was sonicated and spun at the highest speed (21130 g) for 10 minutes and the supernatant was collected and processed for BCA analysis.

*Mouse transmission electron microscopy*

Mice were anesthetized with isoflurane and transcardially perfused with 25 mL of PBS. Hearts and tibialis muscles were rapidly dissected and a 1 mm x 1 mm x 1 mm sample was drop fixed in 4% glutaraldehyde (Electron Microscopy Services, cat# 16020) in electron microscopy (EM) buffer (0.1 M sodium cacodylate at pH 7.4 with 2 mM calcium chloride) for 1 hour at room temperature and then at 4 C for at least 24 hours. Samples were washed with EM buffer and treated with 1% osmium tetroxide in 0.1 M sodium cacodylate buffer at pH 7.4 for 1 hour on ice, washed and *en bloc* stained with 0.25% uranyl acetate in 0.1 M acetate buffer at pH 5.0 overnight at 4 C, dehydrated with a graded series of ethanol washes and finally embedded in epoxy resins. Ultrathin sections (70 nm) were stained with lead citrate and imaged with a JEOL 1200 EXII TEM.

*BN-PAGE*

The mitochondrial fraction was resuspended in 10 mM of HEPES pH 7.6 and 0.5 M sucrose. Protein concentration was measured using the BCA assay. The mitochondrial fraction was solubilized in 1% n-Dodecyl-β-D-Maltoside (DDM) and 1X NativePAGE sample buffer. The NativePAGE Novex Bis-Tris Gel System (Thermo Fisher Scientific) was used according to the manufacturer's instructions with the following modifications: 20 μg of the mitochondrial fraction was loaded, only Light Blue Cathode Buffer was used, and electrophoresis was performed at 150 V for 1 hour then 250 V for 2 hours. Polyvinylidene fluoride (PVDF) was used as the membrane for immunoblotting, and transfer was performed at 130 V for 1 hour at 4 C. The membrane was then washed with 10% acetic acid for 20 minutes and air-dried. Afterwards, the membrane was washed 5 times with methanol to remove residual Coomassie Blue dye, blocked with 5% milk in TBST for 1 hour at room temperature and blotted for proteins of interest.

#### Supplemental Figures

**Figure S1. Related to Figure 1. Phenotypic characterization of the G58R mutation in human and mouse.**

(A) Oil Red O of pectoralis muscle of the proband (III-3) showing significantly increased lipid droplets. (B) Long-range PCR of mtDNA of negative control (NC); proband's left ventricle (LV), right ventricle (RV), and skeletal muscle (pectoralis, SM); and positive control (PC). (C) Histochemistry of proband's mother's (II-2) vastus lateralis showing COX-negative fibers and occasional ragged-red fibers. (D) Sanger sequencing of C10 from a $C10^{G58R}$ mouse showing the heterozygous G58R mutation. (E) Weights of $C10^{WT}$ and $C10^{G58R}$ mice. At least 3 mice per group per timepoint. At least 10 mice per group total. (F) Time to descend a 50cm pole for 18-week-old $C10^{WT}$ and $C10^{G58R}$ mice, and 25-week-old $C10^{S59L}$ mice. (G) Representative COX activity stain of 36-week-old $C10^{WT}$ and $C10^{G58R}$ mouse skeletal muscles, (at least 3 FOV per mouse from 3 mice per group). (H) Quantification of fiber diameter from (G), (n = 60 fibers from 3 mice for $C10^{WT}$ tibialis and $C10^{G58R}$ soleus, and n = 40 fibers from 2 mice for $C10^{WT}$ soleus and $C10^{G58R}$ tibialis). (I) Left: representative images of LipidTOX staining of 30-week-old $C10^{WT}$, $C10^{G58R}$, and $C10^{S59L}$ tibialis muscles on the $OMA1^{+/-}$ background. Right: quantification of left, (n = 3 mice per genotype; 6 FOV per mouse). (J) EKG lead II of a $C10^{G58R}$ mouse showing second-degree atrioventricular block.

### Supplementary Figure 1

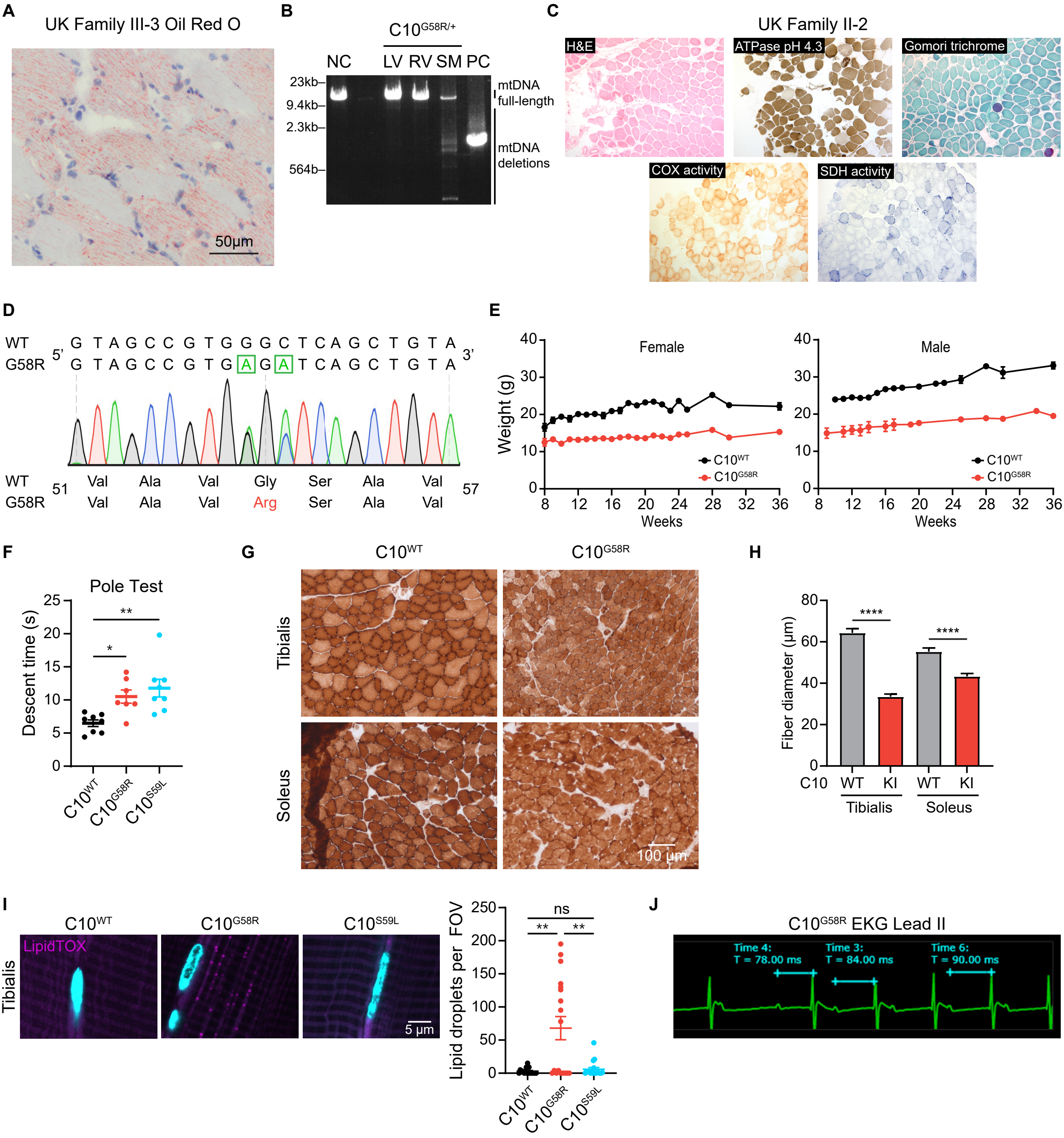

**Figure S2. Related to Figure 1. CLAMS of G58R mice.**

(A) Total food intake over 72 hours, (n = 12 mice per genotype). (B) Absolute (left) and relative (right) lean and fat mass, (n = 12 mice per genotype). (C) Respiratory exchange ratio hourly plot, (n = 12 mice per genotype). (D) Hourly plots (left) and generalized linear model regression (right) of oxygen consumption, carbon dioxide production, and energy expenditure, (n = 12 mice per genotype).

Supplementary Figure 2

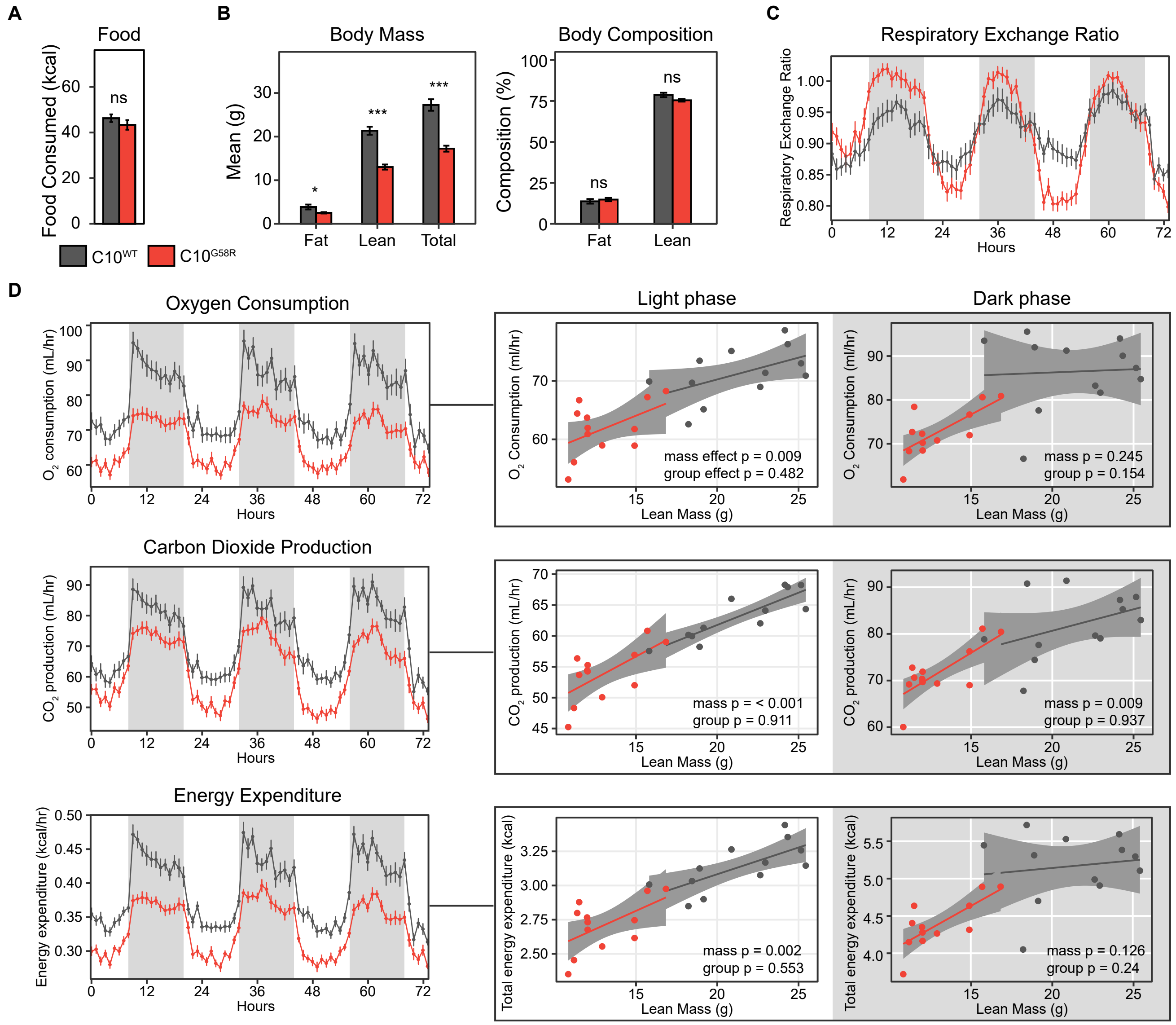

**Figure S3. Related to Figure 2. Effect of C10 mutations on C10 solubility, aggregation, and mitochondrial fragmentation.**

(A) Immunoblots of Triton-X (TX)-soluble and -insoluble C10 from HEK293 C2/C10 DKO lysates after transfection with C10 containing G58 substitutions with amino acids of varying hydrophobicity. (B) Levels of mitochondrial fragmentation in HeLa cells transfected with C10 constructs containing the indicated arginine substitutions in the central α-helix. (C) Left: representative confocal images of staining for C10 in tibialis of 30-week-old C10$^{WT}$, C10$^{G58R}$, and C10$^{S59L}$ mice on the OMA1$^{+/-}$ background. Right: quantification of C10 aggregate area, (n = 3 mice per genotype; 6 FOV per mouse).

Supplementary Figure 3

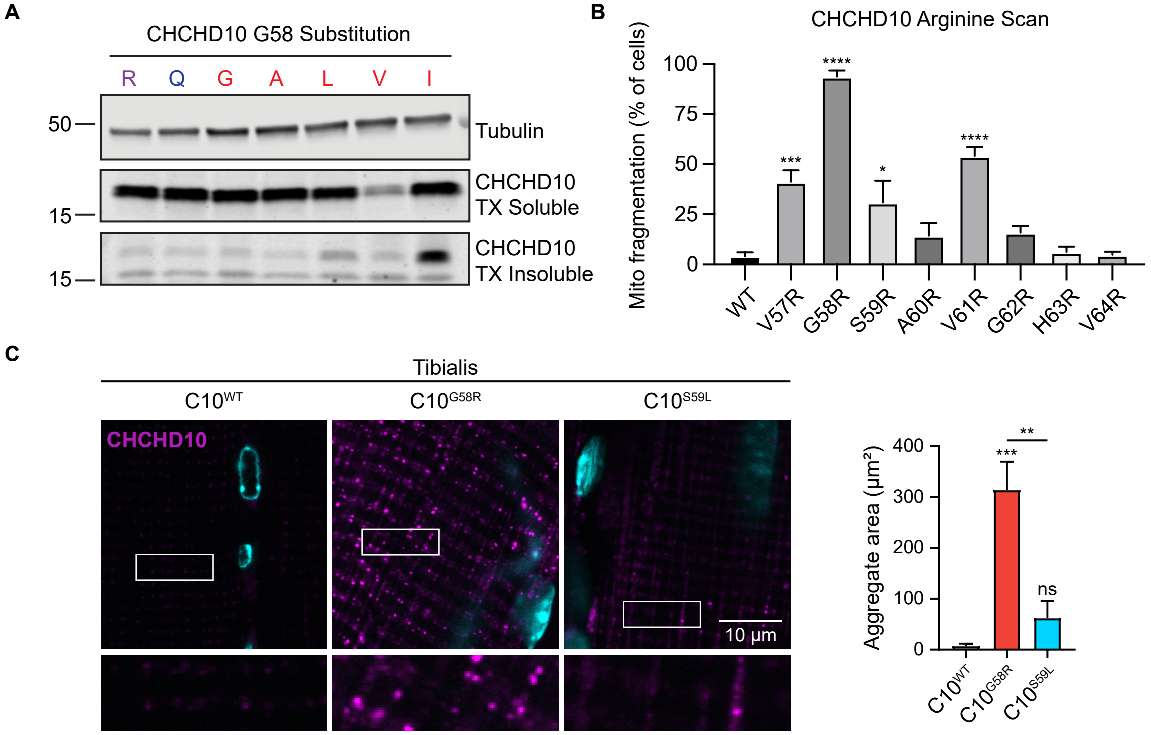

**Figure S4. Related to Figure 4. OMA1 is activated in $C10^{G58R}$ mouse tissues and is required for survival.**

(A) Loading controls for OPA1 and OMA1 (top) and C2/C10 (bottom) in Figure 4A. (B) Confocal images of HEK293 Tet-On C10 G58R OE cells after no treatment, 1 day of doxycycline treatment, and 4 days of doxycycline treatment. (C) Membrane potential estimations from TMRE intensity from HEK293 Tet-On C10 G58R OE treated with DMSO (CTRL) or 1 μg/mL doxycycline overnight, followed by no treatment, 15 min 10 μM CCCP treatment, or 15 min 10 mg/mL oligomycin treatment, (n = 3 biological replicates). (D) Genotypes of pups from the $C10^{WT}$ ; $OMA1^{-/-}$ X $C10^{G58R}$ ; $OMA1^{+/-}$ cross at P1 and P5. (E) Left: representative confocal images showing C10 and p62 staining of 14-week-old mouse hearts. Right: quantification of C10 and p62 aggregate area, (n = 3 mice per genotype for $OMA1^{+/-}$ ; $C10^{WT}$ and $OMA1^{+/-}$ ; $C10^{G58R}$, and n = 4 mice per genotype for and $OMA1^{-/-}$ ; $C10^{WT}$ and $OMA1^{-/-}$ ; $C10^{G58R}$. 10 FOV quantified per mouse). (F) Long-range PCR of a 12.8kb segment of mtDNA from 14-week-old mouse hearts (left) and tibiales (right). DNA ladder is HyperLadder 1kb. Each lane represents an individual mouse. (G) E18.5 mouse embryos from the $C10^{WT}$ ; $OMA1^{-/-}$ X $C10^{G58R}$ ; $OMA1^{+/-}$ cross in the uterus (top) and outside of the uterus (bottom). The $OMA1^{-/-}$ ; $C10^{G58R}$ embryo is indicated. (H) P5 pups from the $C10^{WT}$ ; $OMA1^{-/-}$ X $C10^{G58R}$ ; $OMA1^{+/-}$ cross. (I) P1 and P5 pup weights from the $C10^{WT}$ ; $OMA1^{-/-}$ X $C10^{G58R}$ ; $OMA1^{+/-}$ cross. (J) Expected vs observed numbers of pup genotypes from the $C10^{WT}$ ; $OMA1^{-/-}$ X $C10^{S59L}$ ; $OMA1^{+/-}$ cross.

### Supplementary Figure 4

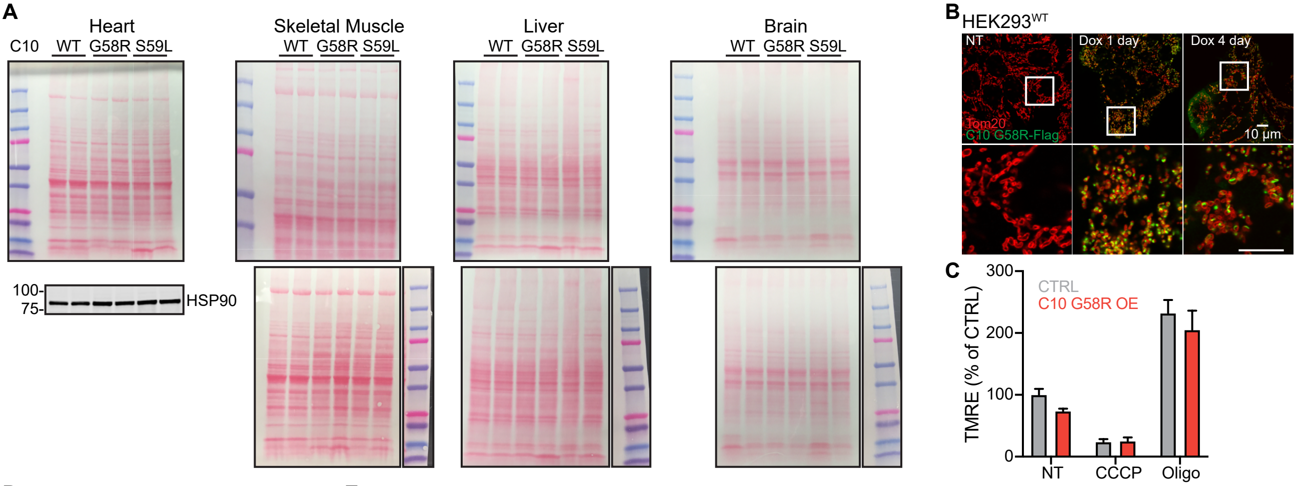


D

Neonatal genotypes

| Genotype | P1 | P5 |
|---|---|---|
| C10$^{WT}$;OMA1$^{+/-}$ | 6 | 6 |
| C10$^{WT}$;OMA1$^{-/-}$ | 6 | 9 |
| C10$^{G58R}$;OMA1$^{+/-}$ | 6 | 5 |
| C10$^{G58R}$;OMA1$^{-/-}$ | 4 | 7 |

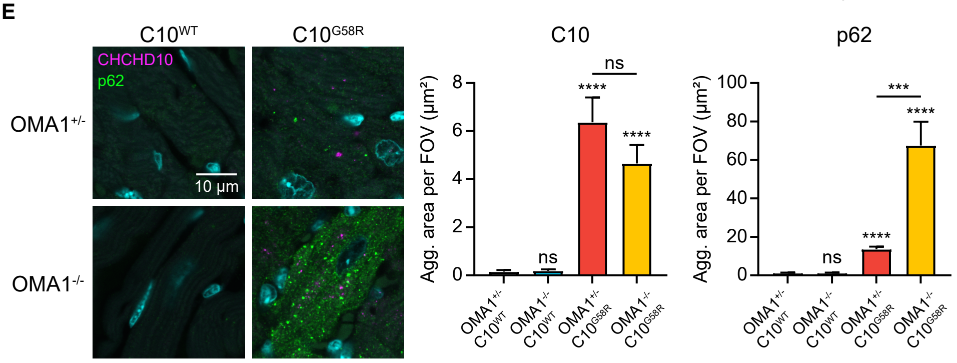


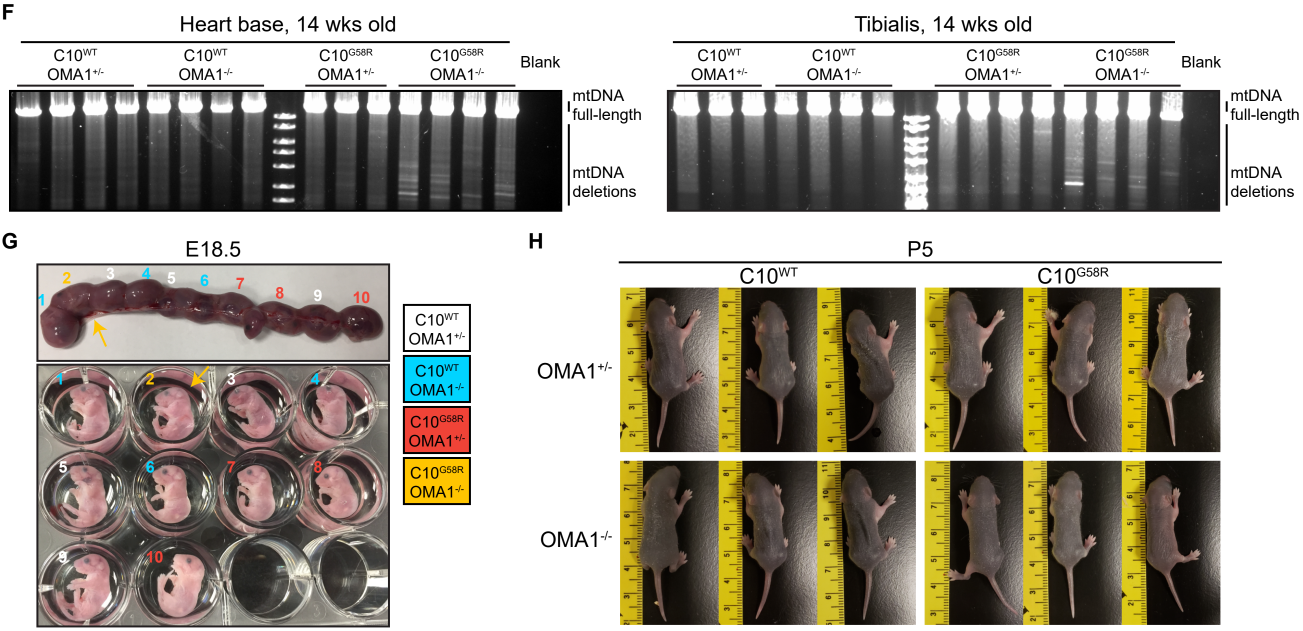


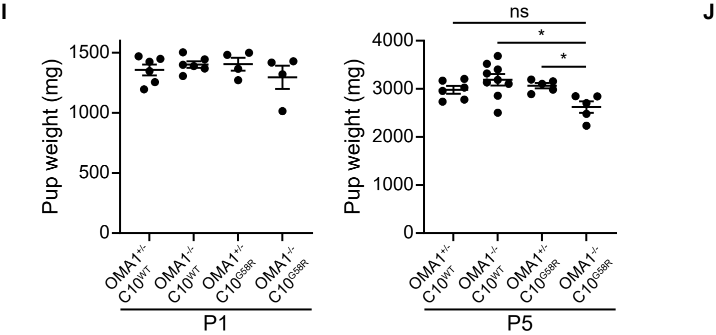


J

Genotypes at weaning

| Genotype | Exp | Obs |
|---|---|---|
| C10$^{WT}$;OMA1$^{+/-}$ | 17 | 18 |
| C10$^{WT}$;OMA1$^{-/-}$ | 17 | 15 |
| C10$^{S59L}$;OMA1$^{+/-}$ | 17 | 21 |
| C10$^{S59L}$;OMA1$^{-/-}$ | 17 | 14 |

$p = 0.6226$

**Figure S5. Related to Figure 5. Representative TEM images of megamitochondria.**

Supplementary Figure 5

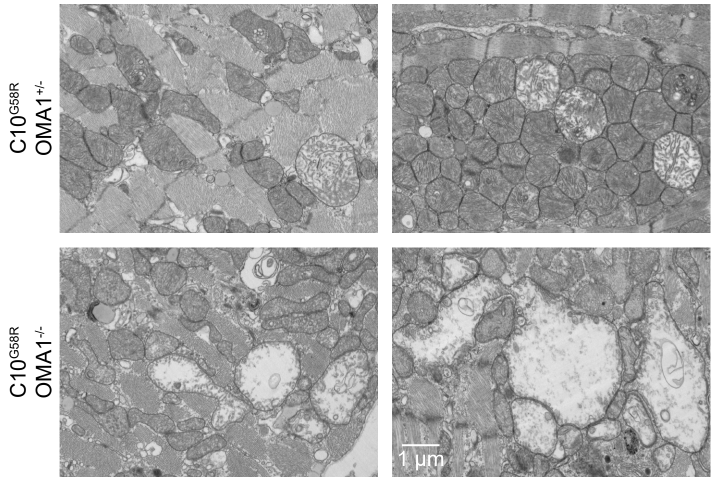

**Figure S6. Related to Figure 6. Effect of OMA1 ASO on mouse health, and effect of OMA1 on transcription.**

(A) Weight of $C10^{G58R}$ ; $OMA1^{+/-}$ mice before the first ASO injection and after the last injection. (B) Survival of $C10^{G58R}$ ; $OMA1^{+/-}$ mice over the course of ASO treatment, (n = 5 mice for CTRL and OMA1 ASO#1, and n = 4 mice for OMA1 ASO#2). (C) Grip strength, treadmill, rotarod, and pole test results of 33-week-old $C10^{G58R}$ ; $OMA1^{+/-}$ mice after 12 weeks of CTRL ASO or OMA1 ASO. (D) Percent ejection fraction and pulmonary artery peak velocity of 33-week-old $C10^{G58R}$ ; $OMA1^{+/-}$ mice after 12 weeks of CTRL ASO or OMA1 ASO. (E) Left: long-range PCR of a 12.8kb segment of mtDNA from hearts of 33-week-old $C10^{G58R}$ ; $OMA1^{+/-}$ mice after 12 weeks of CTRL ASO or OMA1 ASO. Right: mtDNA copy number. (F) Loading controls for Figure 6E. (G) Effect of OMA1 ASO on expression of G58R DEGs identified in the 14-week-old $C10^{G58R}$ ; $OMA1^{+/-}$ vs $C10^{WT}$ ; $OMA1^{+/-}$ comparison. (H) Microarray data of hearts from mice of the indicated genotypes. Each column represents a mouse. (I) Foldchange of the top 10 DEGs of the OMA1 ASO vs CTRL ASO comparison in said comparison, the $C10^{G58R}$ ; $OMA1^{-/-}$ vs $C10^{G58R}$ ; $OMA1^{+/-}$ comparison, and the $C10^{S59L}$ ; $OMA1^{-/-}$ vs $C10^{S59L}$ ; $OMA1^{+/-}$ comparison.

### Supplementary Figure 6

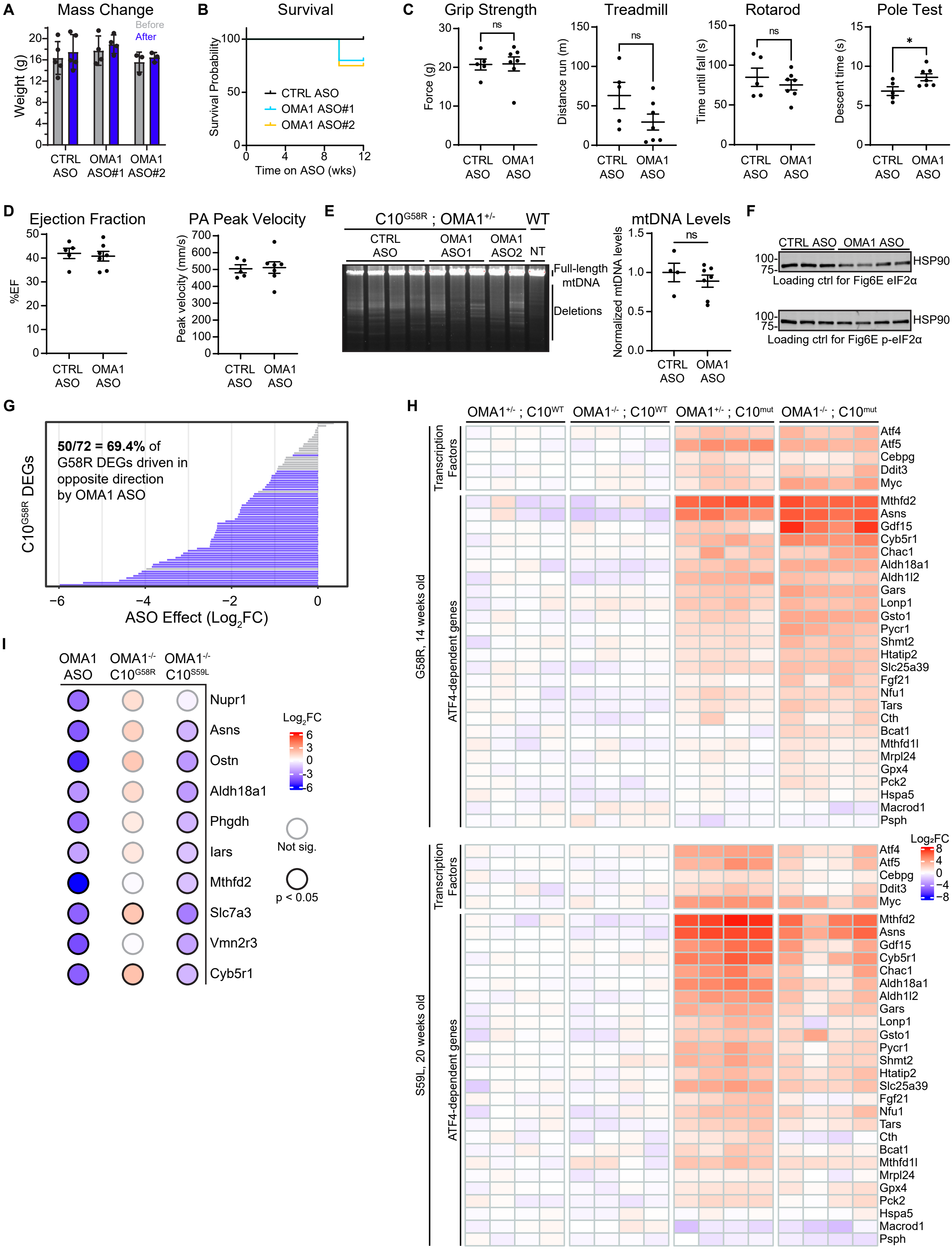

**Figure S7. Related to Figure 7. Effects of different models of mitochondrial stress on the mitochondrial proteome.**

(A) Loading controls for Figure 7B. (B) Loading controls for Figure 7C. (C) BN-PAGE of $C10^{WT}$ and $C10^{G58R}$ heart mitochondrial lysates showing where the gel was cut for downstream proteomics. (D) Fold changes of CIV subunits between $C10^{WT}$ and $C10^{G58R}$ heart mitochondrial lysates in the different BN-PAGE slices from (C). Liver isoforms are in red and heart isoforms in blue. The dashed line indicates the median FC, (n = 3 mice per group). (E) Protein fold changes vs WT of ATF4-dependent proteins, CI-V subunits, and OMA1 in $C10^{G58R}$ and different models of mitochondrial disease, resulting from heart KO of the following genes Twnk, Tfam, Polrmt, Lrpprc, and Mterf4. Data is from Kühl et al., 2017. (F) Fold changes of CIV subunits in models in (E). Data is from Kühl et al., 2017. Liver isoforms are in red and heart isoforms in blue. The dashed line indicates the median FC.

### Supplementary Figure 7

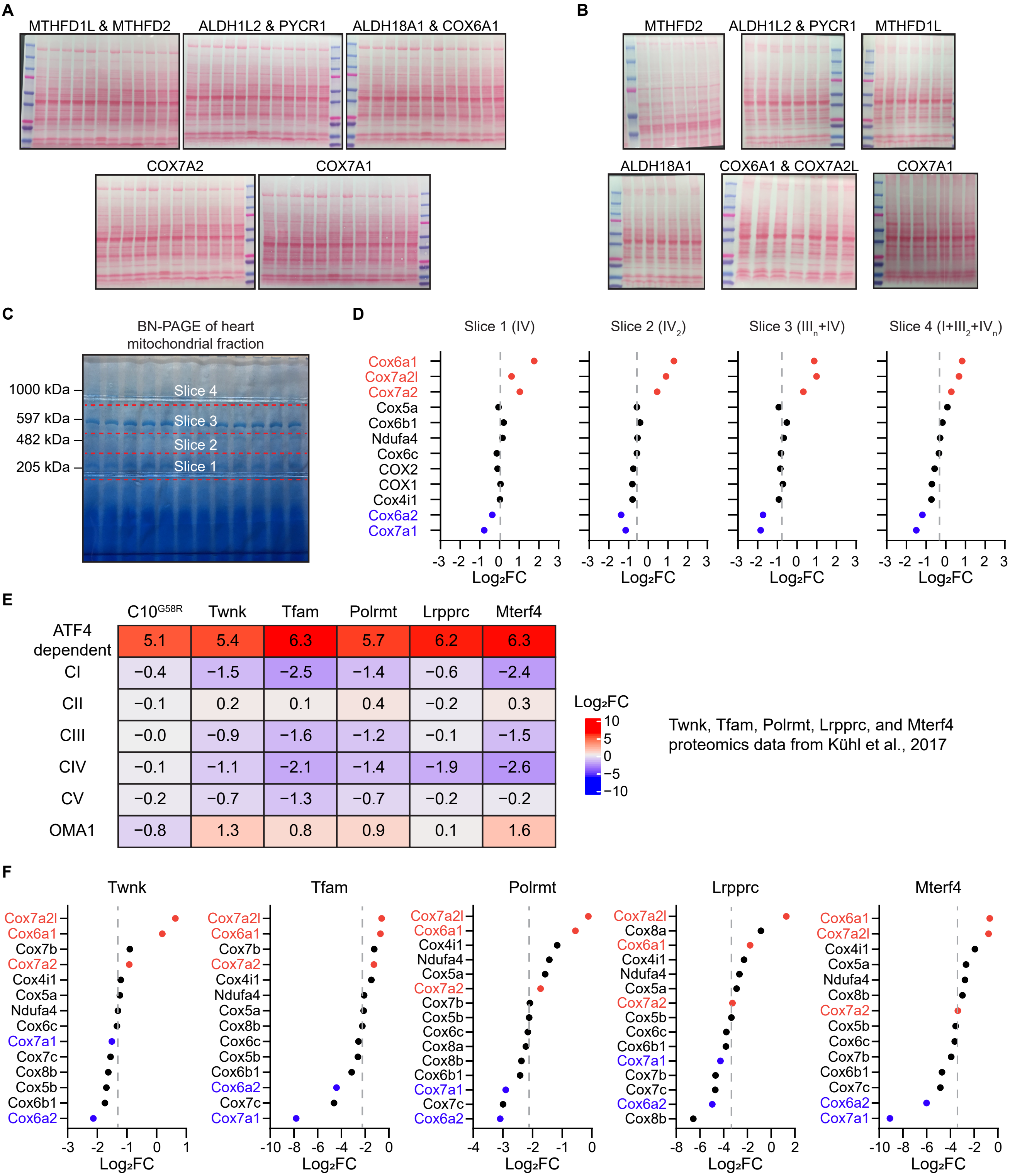
